# Defining the functional significance of intergenic transcribed regions

**DOI:** 10.1101/127282

**Authors:** John P. Lloyd, Zing Tsung-Yeh Tsai, Rosalie P. Sowers, Nicholas L. Panchy, Shin-Han Shiu

## Abstract

With advances in transcript profiling, the presence of transcriptional activities in intergenic regions has been well established. However, whether intergenic expression reflects transcriptional noise or activity of novel genes remains unclear. We identified intergenic transcribed regions (ITRs) in 15 diverse flowering plant species and found that the amount of intergenic expression correlates with genome size, a pattern that could be expected if intergenic expression is largely nonfunctional. To further assess the functionality of ITRs, we first built machine learning classifiers using *Arabidopsis thaliana* as a model that accurately distinguish functional sequences (phenotype genes) and likely nonfunctional ones (pseudogenes and unexpressed intergenic regions) by integrating 93 biochemical, evolutionary, and sequence-structure features. Next, by applying the models genome-wide, we found that 4,427 ITRs (38%) and 796 annotated ncRNAs (44%) had features significantly similar to benchmark protein-coding or RNA genes and thus were likely parts of functional genes. Approximately 60% of ITRs and ncRNAs were more similar to nonfunctional sequences and were likely transcriptional noise. The predictive framework established here provides not only a comprehensive look at how functional, genic sequences are distinct from likely nonfunctional ones, but also a new way to differentiate novel genes from genomic regions with noisy transcriptional activities.

## INTRODUCTION

Advances in sequencing technology have helped to identify pervasive transcription in intergenic regions with no annotated genes. These intergenic transcripts have been found in metazoa and fungi, including human (ENCODE Project Consortium 2012), *Drosophila melanogaster* (Brown et al. 2014), *Caenorhabditis elegans* (Boeck et al. 2016), and *Saccharomyces cerevisiae* (Nagalakshmi et al. 2008). In plants, 7,000 to 15,000 intergenic transcripts have also been reported in *Arabidopsis thaliana* (Yamada et al. 2003; Stolc et al. 2005; Moghe et al. 2013; Krishnakumar et al. 2015) and *Oryza sativa* (Nobuta et al. 2007). The presence of intergenic transcripts indicates that there may be additional genes in genomes that have escaped gene finding efforts thus far, including those that function as RNA genes (Simon and Meyers 2011; Guil and Esteller 2012; Fei et al. 2013; Tan et al. 2015). Meanwhile, it is also possible that some of these intergenic transcripts are products of un-regulated noise (Struhl 2007). Given the functional significance of most intergenic transcripts remains unclear, the identification of functional intergenic transcribed regions (ITRs) represents a fundamental task that is critical to our understanding of genome evolution.

Loss-of-function study represents the gold standard by which the functional significance of genomic regions, including ITRs, can be confirmed (Ponting and Belgard 2010; Niu and Jiang 2013). In *Mus musculus* (mouse), at least 25 ITRs with loss-of-function mutant phenotypes have been identified (Sauvageau et al. 2013; Lai et al. 2015). In human, 162 long intergenic non-coding RNAs harbor phenotype-associated SNPs (Ning et al. 2013). In addition to intergenic expression, most model organisms feature an abundance of annotated non-coding RNA (ncRNA) sequences (Zhao et al. 2016), which are mostly identified through the presence of transcriptional evidence occurring outside of annotated protein-coding genes. Thus, the only difference between ITRs and most ncRNA sequences is whether or not they have been annotated. Similar to the ITR examples above, a small number of ncRNAs have been confirmed as functional through loss-of-function experiments including *Xist* in mouse (Penny et al. 1996; Marahrens et al. 1997), *Malat1* in human (Bernard et al. 2010), *bereft* in *D. melanogaster* (Hardiman et al. 2002), and *At4* in *A. thaliana* (Shin et al. 2006).

However, the number of ITRs and ncRNAs with well-established functions is dwarfed by those without functional evidence. While some ITRs and ncRNAs can be novel genes, intergenic transcription may also be the byproduct of noisy transcription that can occur due to nonspecific landing of RNA Polymerase II (RNA Pol II) or spurious regulatory signals that drive expression in random genomic regions (Struhl 2007). In the ENCODE project (ENCODE Project Consortium 2012), ∼80% of the human genome was defined as biochemically functional as reproducible biochemical activities, e.g. transcription, could be detected. This has drawn considerable critique because the existence of a biochemical activity is not an indication of selection (Eddy 2013; Graur et al. 2013; Niu and Jiang 2013). Instead, it is advocated that a genomic region with discernible activity is only functional if it is under selection (Amundson and Lauder 1994; Graur et al. 2013; Doolittle et al. 2014). Under this “selected effect” functionality definition, ITRs and most annotated ncRNA genes remain functionally ambiguous.

Due to the debate on the definitions of function post-ENCODE, Kellis et al. (Kellis et al. 2014) suggested that evolutionary, biochemical, and genetic evidences provide complementary information to define functional genomic regions. Consistent with this, integration of biochemical and conservation evidence was successful in identifying regions in the human genome that are under selection (Gulko et al. 2014) and classification of human disease genes and pseudogenes (Tsai et al. 2017). In this study, we adopt a similar framework to investigate the functionality of intergenic transcription in plants. We first identified ITRs in 15 flowering plant species with 17-fold genome size differences and evaluated the relationship between the prevalence of intergenic expression and genome size. Next, we established machine learning models using *A. thaliana* data to predict likely-functional ITRs and ncRNAs based on 93 evolutionary, biochemical, and sequence-structure features. Finally, we applied the models to ITRs and annotated ncRNAs to determine whether these functionally ambiguous sequences are more similar to benchmark functional or likely nonfunctional sequences.

## RESULTS & DISCUSSION

### Genome size versus prevalence of intergenic transcripts indicates ITRs may generally be nonfunctional

Transcription of an unannotated, intergenic region could be due to nonfunctional transcriptional noise or the activity of a novel gene. If noisy transcription occurs due to random landing of RNA Pol II or spurious regulatory signals, a naïve expectation is that, as genome size increases, the total nucleotides covered by ITRs would increase accordingly. By contrast, we expect that the extent of expression for genic sequences will not be significantly correlated with 4 genome size because larger plant genomes do not necessarily have more genes (*r*^*2*^=0.01; *p*=0.56).

To gauge if ITRs generally behave more like what we expect of noisy or genic transcription, we first identified genic and intergenic transcribed regions using leaf transcriptome data from 15 flowering plants with 17-fold differences in genome size (Supplementary Table 1). As expected, the coverage of expression originating from annotated genic regions had no significant correlation with genome size (*r*^*2*^=0.03; *p*=0.53; **Fig. 1A**). By contrast, the coverage of intergenic expression was significantly and positively correlated (*r*^*2*^=0.30; *p*=0.04; **Fig. 1B**), consistent with the interpretation that a significant proportion of intergenic expression represents transcriptional noise. However, the correlation between genome size and intergenic expression explained ∼30% of the variation (**Fig. 1B**), suggesting that other factors also affect ITR content, including the possibility that some ITRs are truly functional, novel genes. To further evaluate the functionality of intergenic transcripts, we next identified the biochemical and evolutionary features of functional genic regions and tested whether intergenic transcripts in *A. thaliana* were more similar to functional or nonfunctional sequences.

**Figure 1.**
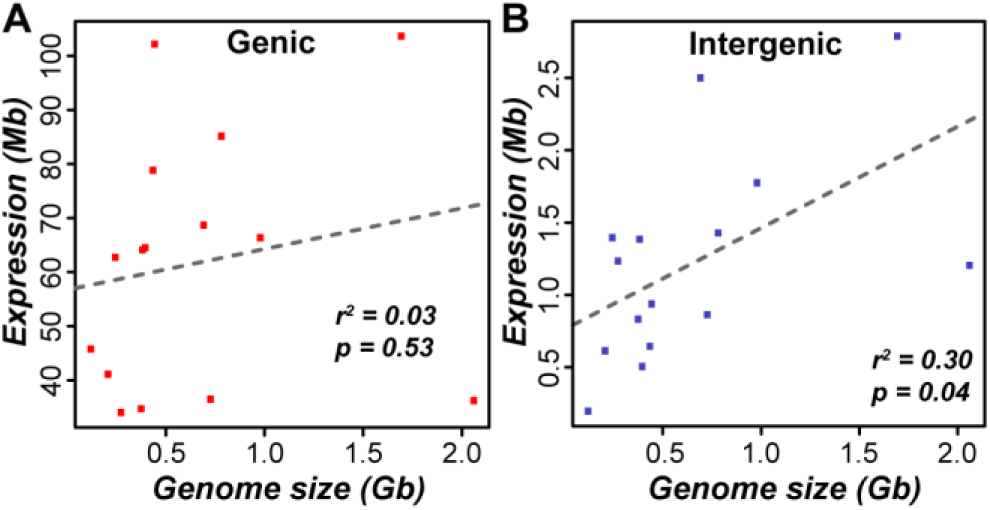
Relationship between genome size and number of nucleotides covered by RNA-seq reads (expression) in 15 flowering plant species. (*A*) annotated genic regions. (*B*) intergenic regions excluding any annotated features. Mb: megabase. Gb: gigabase. Dotted lines: linear model fits, r^2^: square of Pearson’s correlation coefficient.

### Benchmark functional protein-coding and nonfunctional genomic sequences are significantly distinct in multiple features

To determine whether intergenic transcripts resemble functional sequences, we first asked what features allow benchmark functional protein-coding and nonfunctional genomic regions to be distinguished in the model plant *Arabidopsis thaliana*. For benchmark functional sequences, we used protein-coding genes with visible loss-of-function phenotypes when mutated (referred to as phenotype genes, n=1,876; see Materials and Methods). Because their mutations have significant growth and/or developmental impact and likely contribute to reduced fitness, these phenotype genes can be considered functional under the selected effect definition (Neander 1991). For benchmark nonfunctional genomic regions, we utilized pseudogene sequences (n=761; see Materials and Methods). Considering that only 2% of pseudogenes are maintained over 90 million years of divergence between human and mouse (Svensson et al. 2006), it is expected that the majority of pseudogenes are no longer under selection (Li et al. 1981).

We evaluated 93 gene or gene product features for their ability to distinguish between phenotype genes and pseudogenes. These features were grouped into seven categories, including chromatin accessibility, DNA methylation, histone 3 (H3) marks, sequence conservation, sequence-structure, transcription factor (TF) binding, and transcription activity (Supplementary Table 2). We emphasize that no features were exclusive to protein-coding sequences. We used Area Under the Curve - Receiver Operating Characteristic (AUC-ROC) as a metric to measure how well a feature distinguished between phenotype genes and pseudogenes, which ranges between 0.5 (random guessing) and 1 (perfect separation of functional and nonfunctional sequences). Among the seven feature categories, transcription activity features were highly informative (median AUC-ROC=0.88; **Fig. 2A**). Despite the strong performance of transcription activity-related features, the presence of expression (i.e. transcript evidence) was a poor predictor of functionality (AUC-ROC=0.58; **Fig. 2A**). This is because 80% of pseudogenes were considered expressed in ≥1 of 51 RNA-seq datasets, demonstrating that presence of transcripts should not be used by itself as evidence of functionality. Sequence conservation, DNA methylation, TF binding, and H3 mark features were also fairly distinct between phenotype genes and pseudogenes (median AUC-ROC ∼0.7 for each category; **Fig. 2B-E**). By contrast, chromatin accessibility and sequence-structure features were largely uninformative (median AUC-ROC=0.51 and 0.55, respectively; **Fig. 2F,G**). We also observed high performance variability within feature categories (see Supplementary Information). While many features are distinct between phenotype genes and pseudogenes, functional predictions based on single features yield high error rates (Supplementary Table 3; Supplementary Information), indicating a need to jointly consider multiple features for distinguishing phenotype genes and pseudogenes.

**Figure 2.**
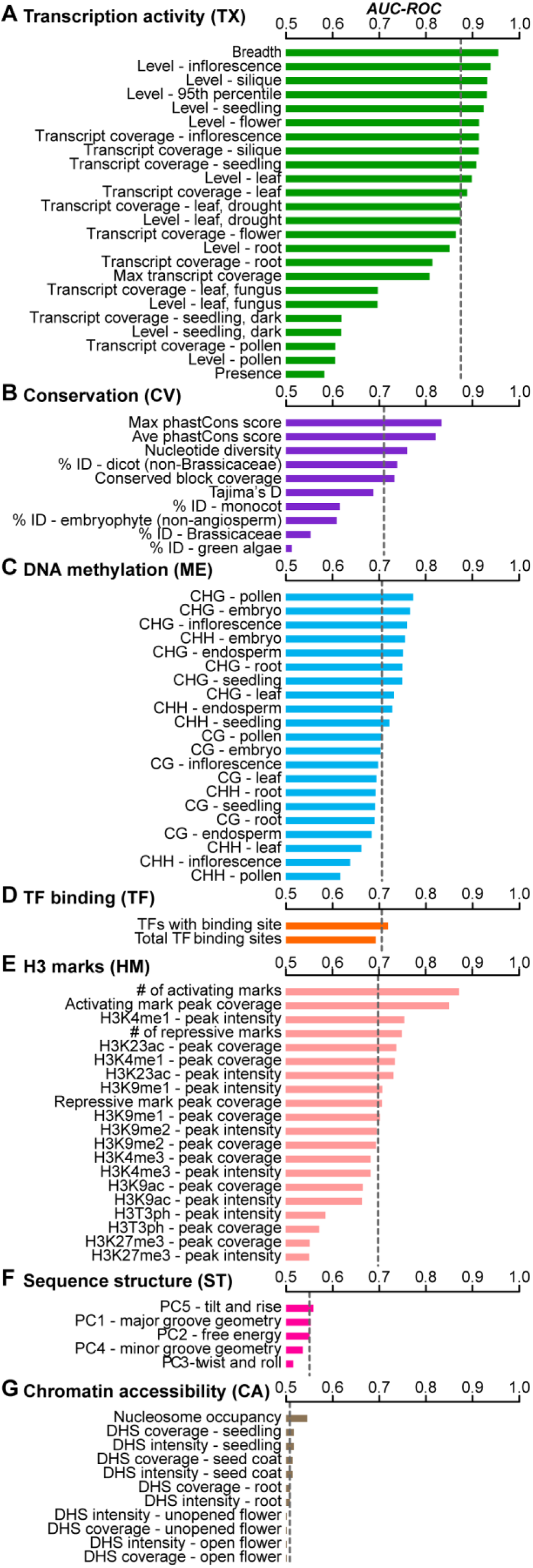
Predictions of functional (phenotype gene) and non-functional (pseudogene) sequences based on each individual feature. Prediction performance is measured using Area Under the Curve-Receiver Operating Characteristic (AUC-ROC). Features include those in the categories of (*A*) transcription activity, (*B*) sequence conservation, (*C*) DNA methylation, (*D*) transcription factor (TF) binding, (*E*) histone 3 (H3) marks, (*F*) sequence structure, and (*G*) chromatin accessibility. AUC-ROC ranges in value from 0.5 (equivalent to random guessing) to 1 (perfect predictions). Dotted lines: median AUC-ROC of features in a category.

### Consideration of multiple features produces accurate predictions of functional genomic regions

To consider multiple features in combination, we instead integrated all 93 features to establish a machine learning model distinguishing phenotype gene and pseudogenes (referred to as the full model; see Materials and Methods). The full model provided more accurate predictions (AUC-ROC=0.98; False Negative Rate (FNR)=4%; False Positive Rate (FPR)=10%; **Fig. 3A**) than any individual feature (**Fig. 2**; Supplementary Fig. 1, Supplementary Table 3). An alternative measure of performance based on the precision (proportion of predicted functional sequences that are truly functional) and recall (proportion of truly functional sequences predicted correctly) values also indicated that the model was performing well (**Fig. 3B**). When compared to the best-performing single feature (expression breadth), the full model had a similar FNR but half the FPR (10% compared to 21%). Thus, the full model is highly capable of distinguishing between phenotype genes and pseudogenes.

**Figure 3.**
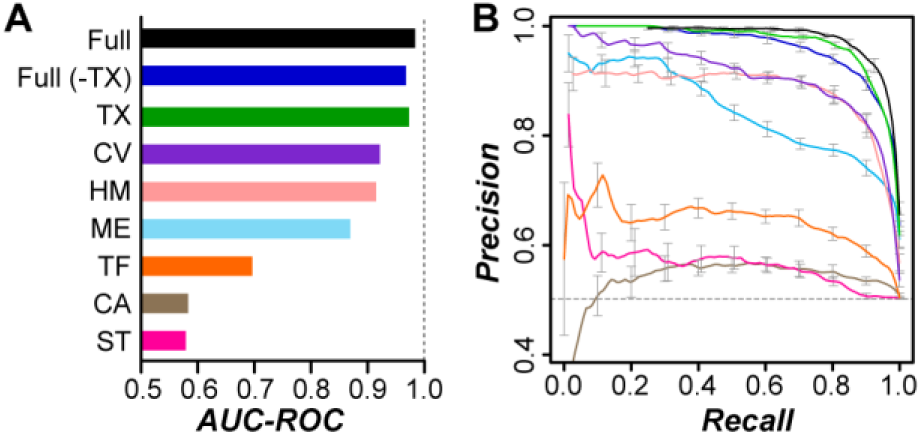
Predictions of functional and nonfunctional sequences based on multiple features. (***A***) AUC-ROC values of function prediction models built when considering all features (Full), all except transcription activity (TX)-related features (Full (-TX)), and all features from each category. The category abbreviations follow those in **Fig. 2**. (***B***) Precision-recall curves of the models with matching colors from (***A***). The models were built using feature values calculated from 500 bp sequence windows. We also conducted principle component (PC) analysis to investigate how well phenotype genes and pseudogenes could be separated and found that phenotype genes (Supplementary Fig. 1A) and pseudogenes (Supplementary Fig. 1B) were distributed in largely distinct space. However, there remained substantial overlap, indicating that standard parametric approaches are not well suited to distinguishing between benchmark functional and nonfunctional sequences.

**Supplementary Figure 1.**
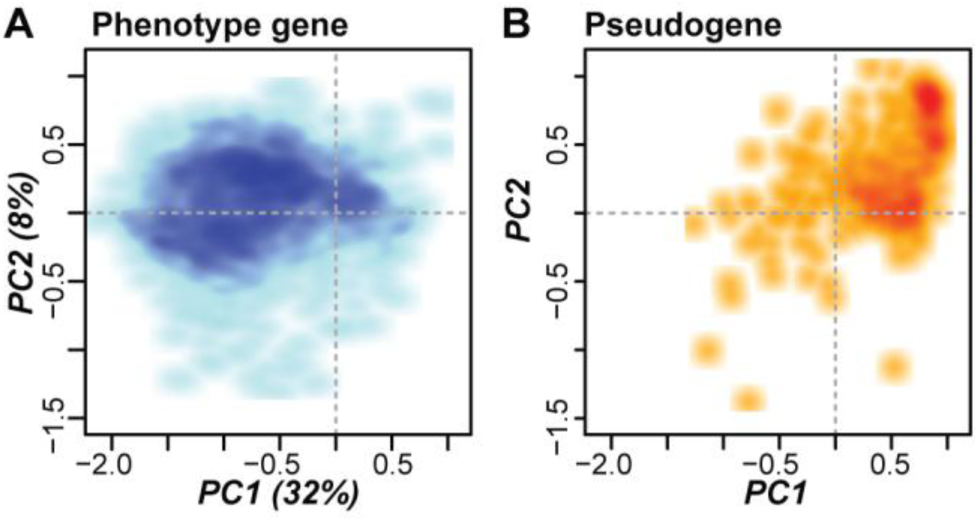
Smoothed scatterplots of the first two principle components (PCs) of (*A*) phenotype gene and (*B*) pseudogene features. The percentages on the axes in *(A)* indicate the feature value variation explained by the associated PC.

We next determined the relative contributions of different feature categories in predicting phenotype genes and pseudogenes and established seven prediction models each using only the subset of features from a single category (**Fig. 2**). Although none of these category-specific models had performance as high as the full model (**Fig. 3C)**, the transcription activity feature category model performed almost as well as the full model (AUC-ROC=0.97, FNR=6%, FPR=12%). Instead of the presence of expression evidence, the breadth and level of transcription are the causes of the strong performance of the transcription activity-only model. We also found that a model excluding transcription activity features (full (-TX), **Fig. 3C,D**) performed almost as well as the full model and similarly to the transcription activity-feature-only model, but with an increased FPR (AUC-ROC=0.96; FNR=3%; FPR=20%). These findings indicate that a diverse array of features can be considered jointly to make highly accurate predictions of the functionality of a genomic sequence. Meanwhile, our finding of the high performance of the transcription activity-only model highlights the possibility of establishing an accurate model for functional prediction in species with only a modest amount of transcriptome data.

### The functional likelihood measure can be used to classify functional and non-functional sequences

To provide a measure of the potential functionality of any sequence in the *A. thaliana* genome, including ITRs and ncRNAs, we utilized the confidence score from the full model as a “functional likelihood” value (Tsai et al. 2017). The functional likelihood (FL) score ranges between 0 and 1, with high values indicating that a sequence is more similar to phenotype genes (functional) and low values indicating a sequence more closely resembles pseudogenes (nonfunctional). FL values for all genomic regions examined in this study are available in Supplementary Table 4. As expected, phenotype genes had high FL values (median=0.97; **Fig. 4A**) and pseudogenes had low values (median=0.01; **Fig. 4B**). To call sequences as functional or not, we defined a threshold FL value (0.35) by maximizing the F-measure (see Materials and Methods). Using this threshold, 96% of phenotype genes (**Fig. 4A**) and 90% of pseudogenes (**Fig. 4B**) are correctly classified as functional and nonfunctional, respectively, demonstrating that the full model is highly capable of distinguishing functional and nonfunctional sequences.

**Figure 4.**
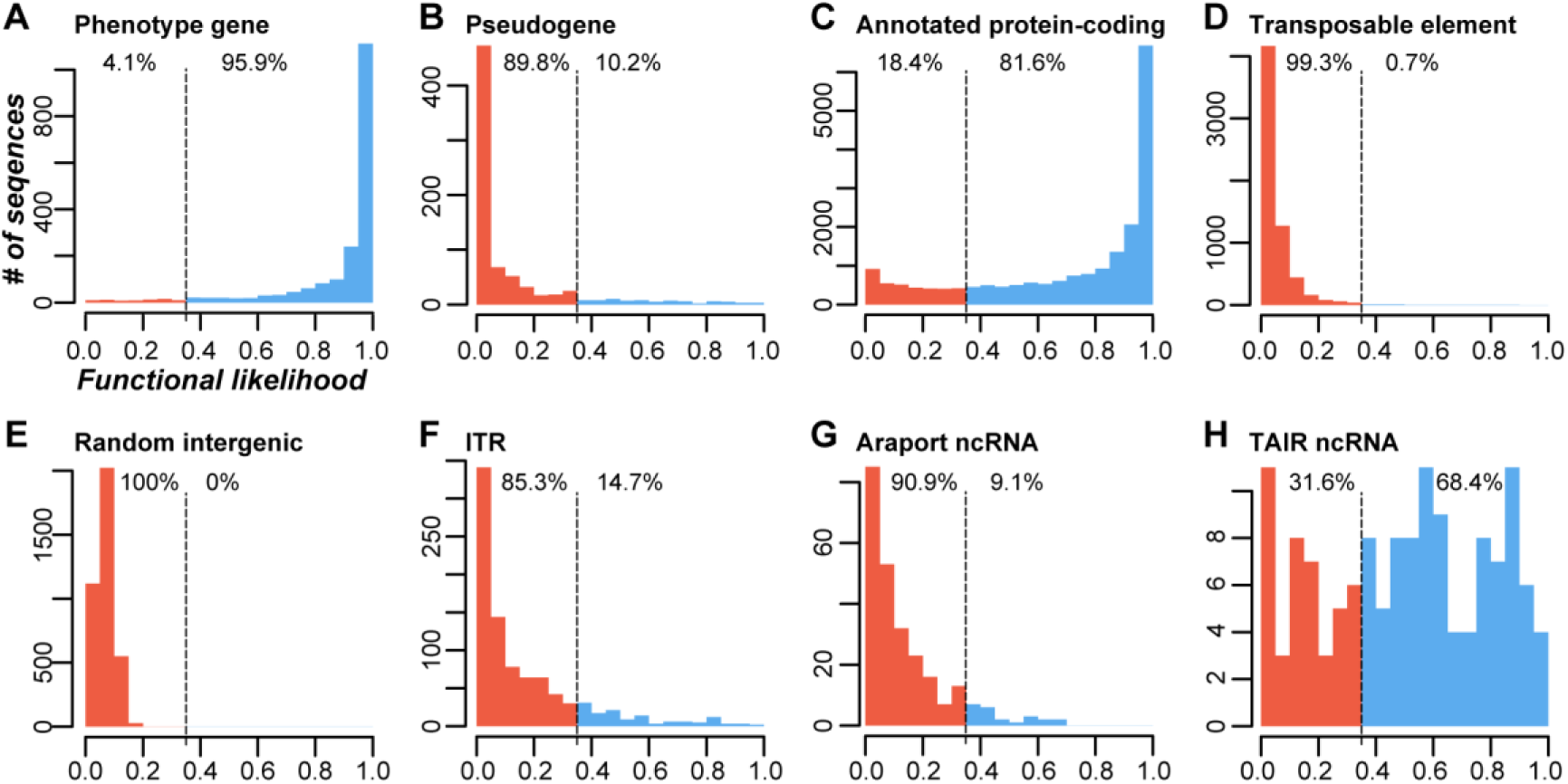
Functional likelihood distributions of various sequence classes based on the full model. (***A***) Phenotype genes. (***B***) Pseudogenes. (***C***) Annotated protein-coding genes. (***D***) Transposable elements. (***E***) Random unexpressed intergenic sequences. (***F***) Intergenic transcribed regions (ITR). (***G***) Araport11 ncRNAs. (***H***) TAIR10 ncRNAs. The full model was established using 500 bp sequence windows. Higher and lower functional likelihood values indicate greater similarity to phenotype genes and pseudogenes, respectively. Vertical dashed lines indicate the threshold for calling a sequence as functional or nonfunctional. The percentages to the left and right of the dashed line indicate the percent of sequences predicted as functional or nonfunctional, respectively.

We next applied our model to predict the functionality of annotated protein-coding genes, transposable elements (TEs), and unexpressed intergenic regions. Most annotated protein-coding genes not included in the phenotype gene dataset had high FL scores (median=0.86; **Fig. 4C**) and 80% were predicted as functional. The features exhibited by low-scoring protein-coding genes and high-scoring pseudogenes are discussed in the Supplementary Information. By contrast, the FLs were low for both TEs (median=0.03, **Fig. 4D**) and unexpressed intergenic regions (median=0.07; **Fig. 4E**), and 99% of TEs and all unexpressed intergenic sequences were predicted as nonfunctional. Overall, the FL measure provides a useful metric to distinguish between phenotype genes and pseudogenes. In addition, the FLs of annotated protein-coding genes, TEs, and unexpressed intergenic sequences agree with *a priori* expectations regarding the functionality of these sequences.

### Most ITRs and annotated ncRNAs do not resemble benchmark phenotype genes

We next evaluated functional predictions of ITRs and ncRNAs. Consistent with previous studies (Moghe et al. 2013), ITRs and ncRNAs in our dataset were more narrowly and weakly expressed and less conserved compared to phenotype genes (Supplementary Fig. 2A,B). In addition, ITRs in particular had biochemical characteristics that were generally more similar to pseudogenes (Supplementary Fig. 2C-F). Given the association between transcription activity features and functional predictions (**Fig 2A; Fig. 3A)**, we investigated how functional prediction models performed for conditionally-functional and narrowly-expressed sequences before applying them to ITRs and ncRNAs. We found that genes with conditional phenotypes had no significant differences in FLs (median=0.96) as those with phenotypes under standard growth conditions (median=0.97; U test, *p*=0.38, Supplementary Fig. 3A), indicating the full model can capture conditionally functional sequences. However, the full model is biased against narrowly-expressed (≤3 tissues) phenotype genes as 65% of them were predicted as nonfunctional (Supplementary Fig. 3B). Further, pseudogenes that were more highly and broadly expressed were disproportionately predicted as functional (**Fig. 5**; Supplementary Fig. 3B). To tailor functional predictions to narrowly-expressed sequences, particularly ITRs and ncRNAs, we generated a “tissue-agnostic” model by excluding expression breadth and features available across multiple tissues (see Materials and Methods). This tissue-agnostic model performed similarly to the full model (AUC-ROC=0.97; FNR=4%; FPR=15%; Supplementary Fig. 4; Supplementary Table 4), although there was a 5% increase in FPR (from 10% to 15%). Importantly, the proportion of phenotype genes expressed in ≤3 tissues predicted as functional increased by 23% (35% in the full model to 58% in the tissue-agnostic model; Supplementary Fig. 3C), indicating that the tissue-agnostic model is more suitable for predicting the functionality of narrowly-expressed sequences than the full model.

**Figure 5.**
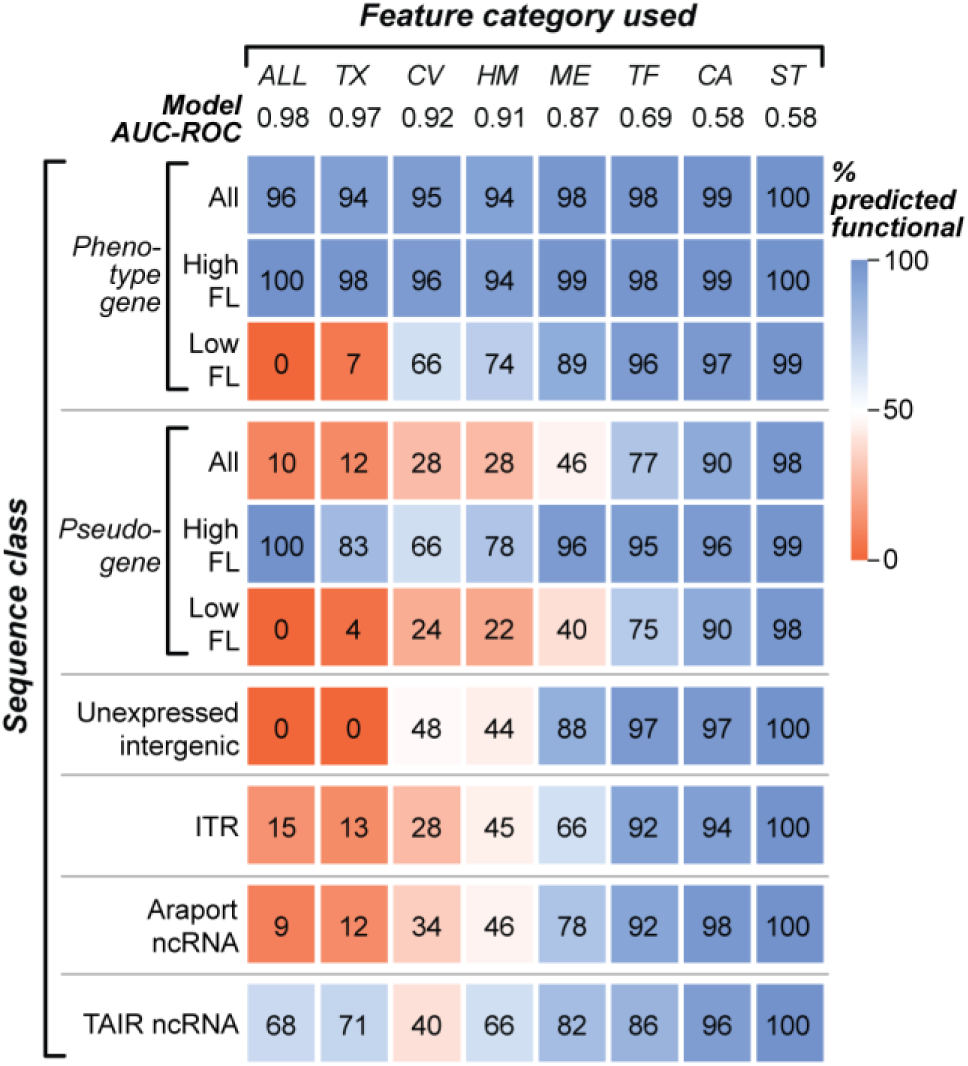
Proportion of phenotype genes, pseudogenes, ITRs, and ncRNAs predicted as functional in the full and single-category models. Percentages of sequence classes that are predicted as functional in models based on all features and the single category models, each using all features from a category (abbreviated according to **Fig. 2**). The models are sorted from left to right based on performance (AUC-ROC). The colors of and numbers within the blocks indicate the proportion sequences predicted as functional by a given model. Phenotype gene and pseudogene sequences are shown in three sub-groups: all sequences (All), and those predicted as functional (high functional likelihood (FL)) and nonfunctional (low FL) in the full model. ITR: intergenic transcribed regions. A greater proportion of ITRs and Araport ncRNAs are predicted as functional when considering only DNA methylation or H3 mark features compared to the full (**Fig. 3**) or tissue-agnostic (Supplementary Fig. 4) models. However, these two category-specific models also had higher false positive rates (unexpressed intergenic sequences and pseudogenes). Thus, these single feature-category models do not provide additional support for the functionality of most Araport ncRNAs and ITRs.

**Supplementary Figure 2.**
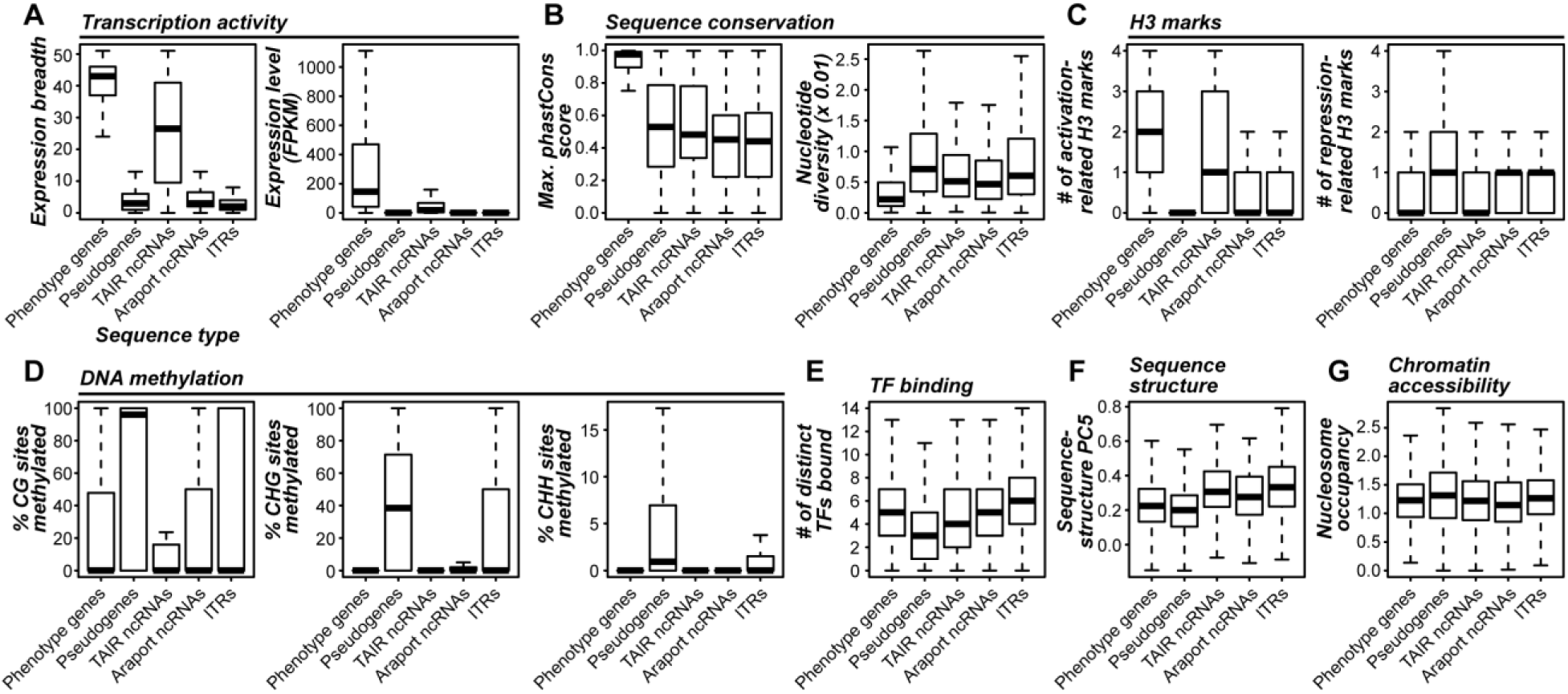
Distributions of 12 example features from (***A***) transcription activity, (***B***) sequence conservation, (***C***) histone 3 (H3) marks, (***D***) DNA methylation, (***E***) transcription factor (TF) binding, (***F***) sequence structure, and (***G***) chromatin accessibility feature categories (Fig. 2). Feature distributions are shown for phenotype genes, pseudogenes, TAIR- and Araport-annotated ncRNAs and intergenic transcribed regions (ITR).

**Supplementary Figure 3.**
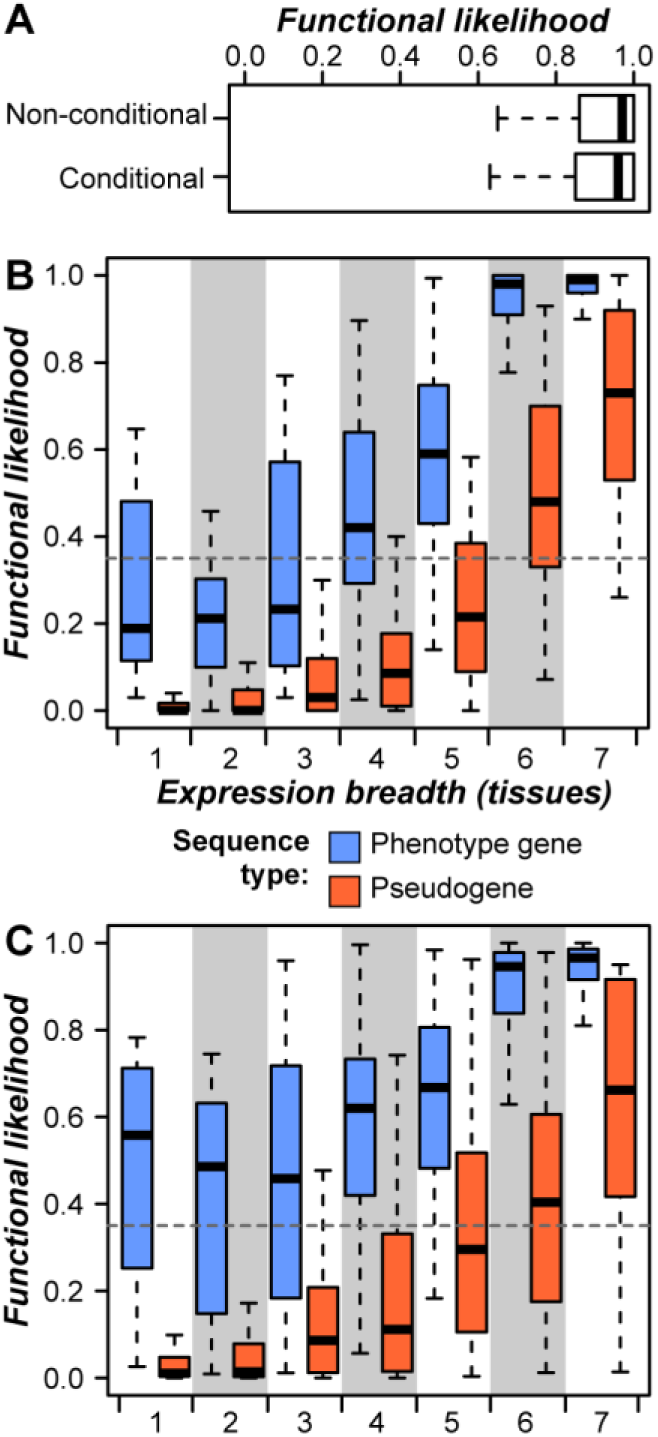
Impacts of conditional phenotypes and expression breadth on the function prediction model. (*A*) Functional likelihood distributions of phenotype genes with mutant phenotypes under standard growth conditions (non-conditional) and non-standard growth conditions such as stressful environments (conditional) based on the 500 bp full model. Feature values were calculated from a random 500 bp region from within the sequence body. Higher and lower functional likelihood values indicate a greater similarity to phenotype genes and pseudogenes, respectively. (*B*,*C*) Distributions of functional likelihood scores for phenotype genes (blue) and pseudogenes (red) for sequences with various breadths of expression for (*B*) the 500 bp full model and (*C*) the 500 bp tissue-agnostic model generated by excluding the expression breadth and features available from multiple tissues. The tissue-agnostic model is aimed toward minimizing the effects of biochemical activity occurring across multiple tissues and predicts a greater proportion of narrowly-expressed phenotype genes as functional compared to the full model.

**Supplementary Figure 4.**
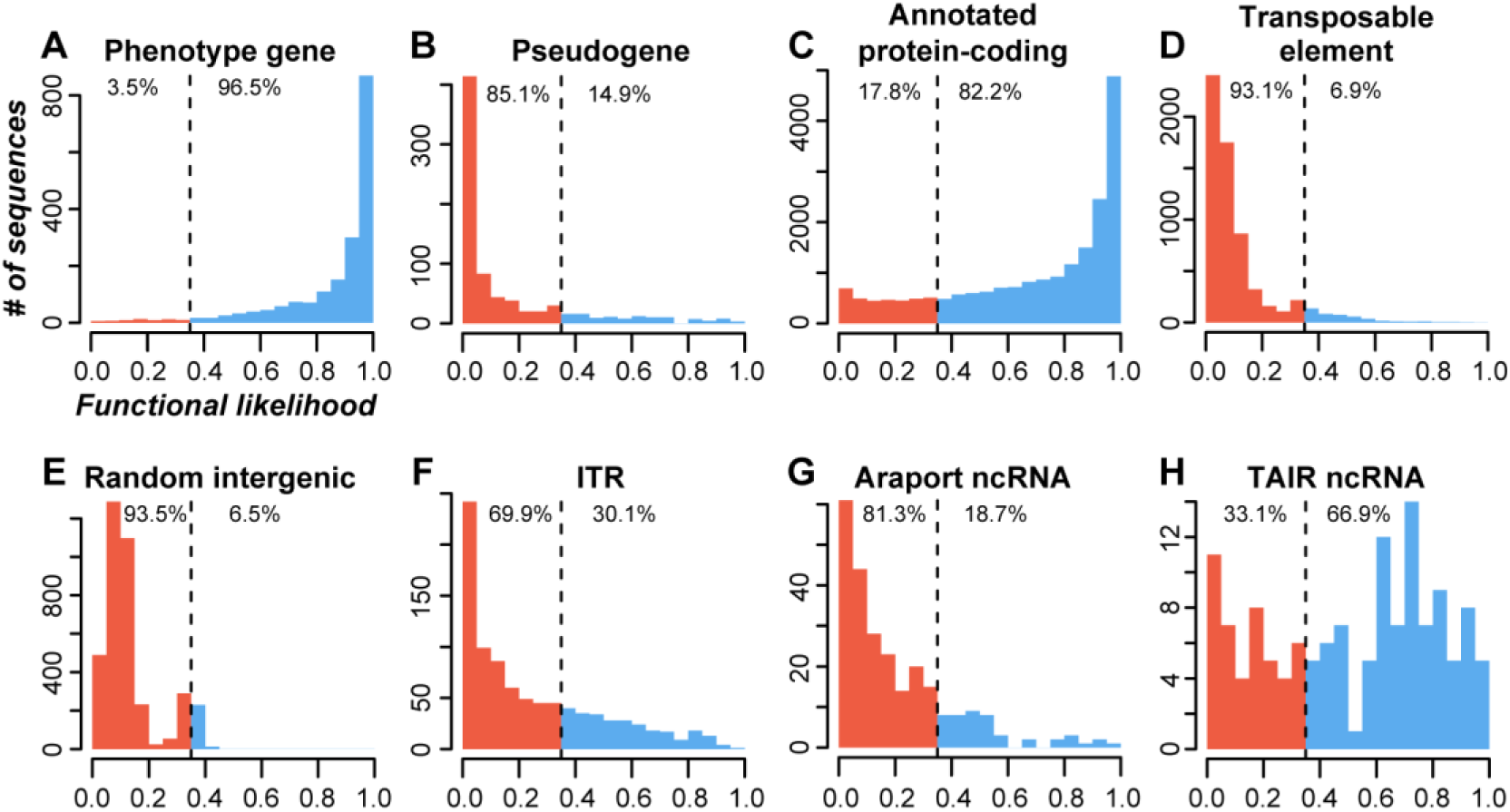
Distributions of functional likelihood scores based on the 500 bp tissue-agnostic model. (*A*) Phenotype genes. (*B*) Pseudogenes. (*C*) Annotated protein-coding genes. (*D*) Transposable elements. (*E*) Random unexpressed intergenic sequences. (*F*) Intergenic transcribed regions (ITR). (*G*) Araport11 ncRNAs. (*H*) TAIR10 ncRNAs. Vertical dashed lines display the threshold to define a sequence as functional or nonfunctional. The numbers to the left and right of the dashed line show the percentage of sequences predicted as functional or nonfunctional, respectively.

We next applied both the full and tissue-agnostic models to 895 ITRs, 136 ncRNAs annotated by The Arabidopsis Information Resource (TAIR), and 252 long ncRNAs annotated by the Araport database that do not overlap with any other annotated genome features. The median FLs based on the full model were low (0.09) for both ITRs (**Fig. 4F**) and Araport ncRNAs (**Fig. 4G**), and only 15% and 9% of these sequences were predicted as functional, respectively. By contrast, TAIR ncRNAs had a significantly higher median FL value (0.53; U tests, both *p*<5e-31; **Fig. 4H**) and 68% were predicted as functional, which is best explained by differences in features from the transcription activity category (**Fig. 5**). We also note that ITRs and annotated ncRNAs that were close to genes were more frequently predicted as functional, suggesting a subset may represent unannotated exons of known genes (Supplementary Fig. 5; Supplementary Information). Next we applied the tissue-agnostic model to ITRs and TAIR/Araport ncRNAs. Compared to the full model, around twice as many ITRs (30%) and Araport ncRNAs (19%) but a similar number of TAIR ncRNA (67%) were predicted as functional. Considering the union of the full and tissue-agnostic model predictions, 268 ITRs (32%), 57 Araport ncRNAs (23%), and 105 TAIR ncRNAs (77%) were likely functional. Thus, the majority of ITRs and Araport ncRNAs are more similar to pseudogenes than to phenotype genes that are predominantly protein coding.

**Supplementary Figure 5.**
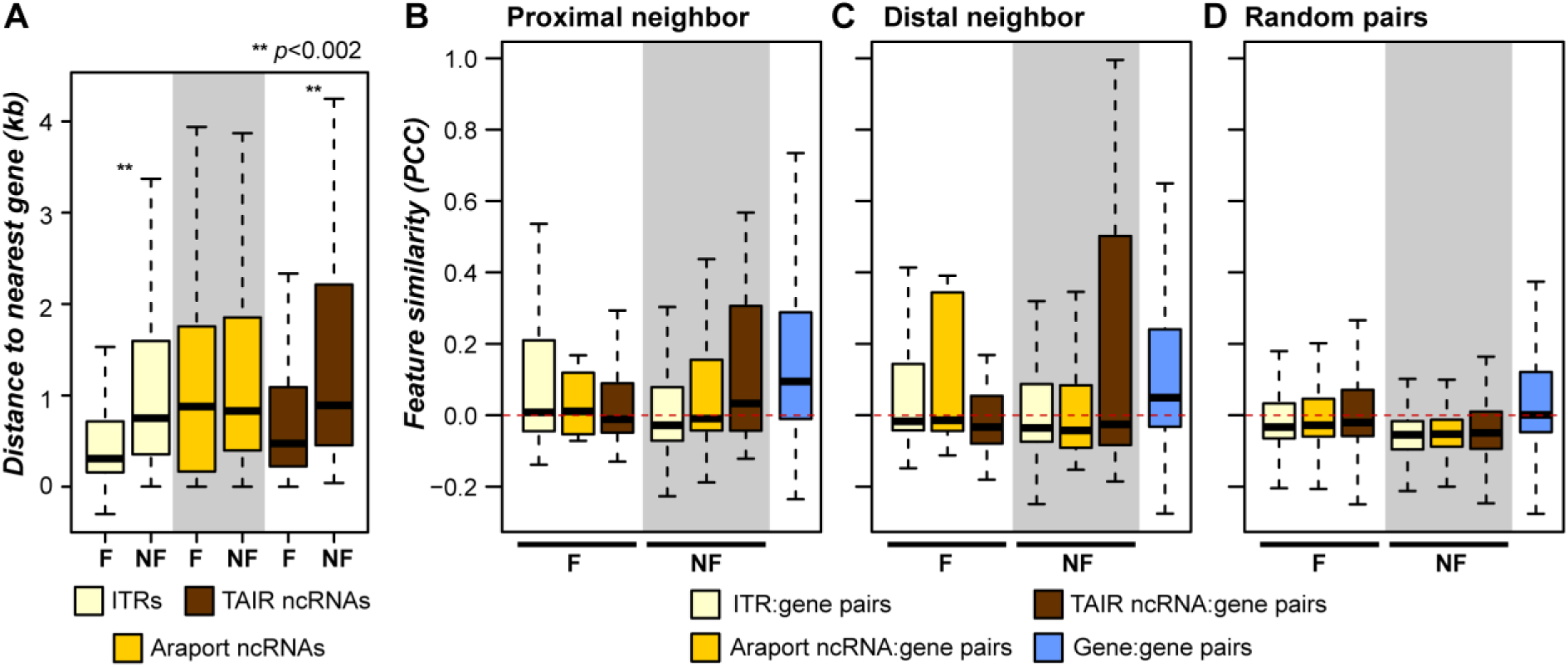
Distance of ITRs and annotated ncRNA regions to and feature similarity with neighboring genes. (*A*) Distance from intergenic transcribed regions (ITRs) and annotated ncRNAs to the closest neighboring gene. ITR and ncRNA sequences are separated by whether they are predicted as functional (F) or nonfunctional (NF) by the 500 bp full model. (*B*) Feature similarity between proximal neighbors (within 95th percentile (456 bp) of intron lengths), and (*C*) distal neighbors (>456 bp). Pairs involving ITRs and annotated ncRNAs were divided by whether the ITR or ncRNA sequence was predicted as functional (F) or nonfunctional (NF) by the full model. Feature values were quantile normalized prior to calculating correlations. (*D*) Feature similarity based on Pearson’s Correlation Coefficients (PCC) between random pairs of ITRs, Araport11 ncRNAs, TAIR10 ncRNAs, or annotated genes.

### Benchmark protein-coding and RNA genes exhibit distinct characteristics

We demonstrated that the majority of ITR and annotated ncRNA sequences do not exhibit characteristics of benchmark phenotype genes. Note that the phenotype genes are predominantly protein coding. Although the features utilized to generate functional predictions were not exclusive to protein-coding sequences, RNA genes may exhibit a distinct feature profile from protein-coding genes. The full and tissue agnostic models described above were established with 500 bp windows and most known RNA genes are too short to be considered by these models. Thus, to evaluate functional predictions among annotated RNA genes, we generated a new tissue-agnostic model using 100 bp sequences (for features, see Supplementary Table 6) that performed similarly to the full 500 bp model, except with 9% higher FNR (AUC-ROC=0.97; FNR=13%; FPR=5%; Supplementary Fig. 6). With this new tissue agnostic model, 50% (three out of six) of RNA genes with documented mutant phenotypes (phenotype RNA genes) were predicted as functional (Supplementary Fig. 6I). We also applied this model to other RNA Pol II-transcribed RNA genes (without documented phenotypes) and found that 15% of microRNA (miRNA) primary transcripts (Supplementary Fig. 6J), 73% of small nucleolar RNAs (snoRNAs; Supplementary Fig. 6K), and 50% of small nuclear RNAs (snRNAs; Supplementary Fig. 6L) were predicted as functional. Although the proportion of phenotype RNA genes predicted as functional (50%) is significantly higher than the proportion of pseudogenes predicted as functional (5%, FET, *p* < 0.004), this finding suggests that a model trained using protein-coding genes has a substantial FNR for detecting RNA genes.

**Figure 6.**
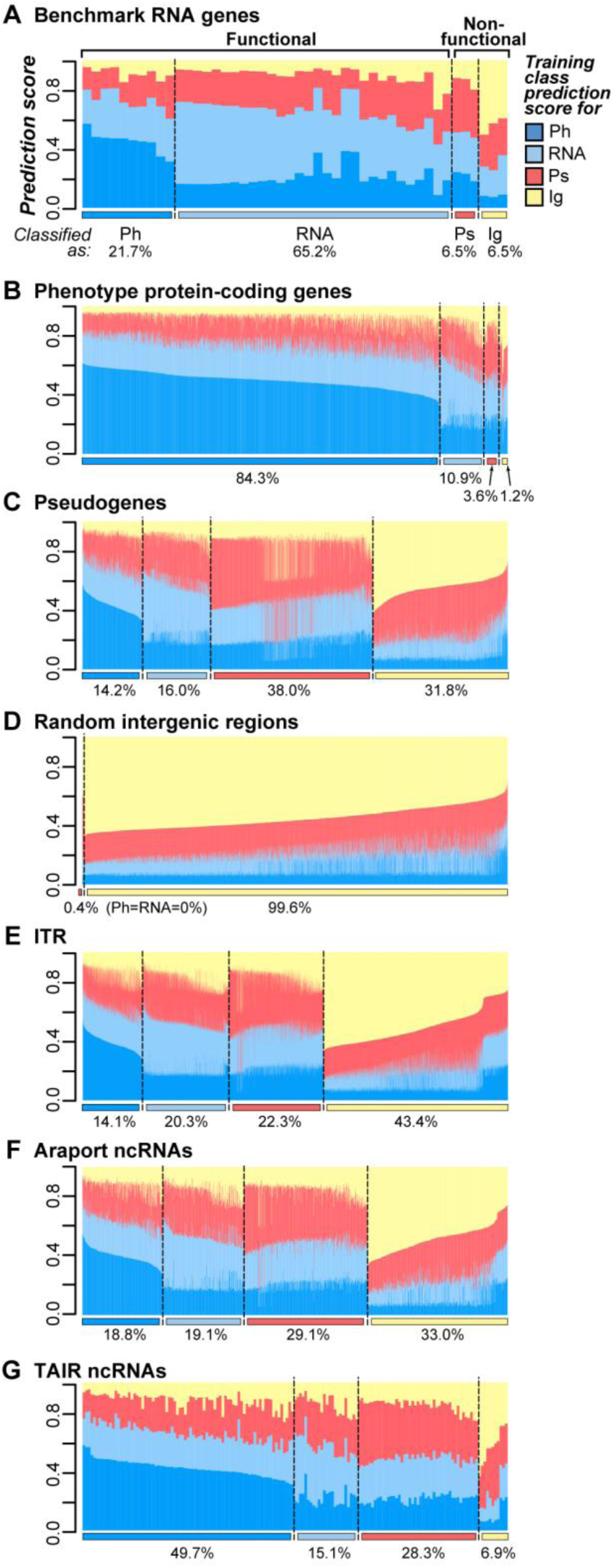
Function predictions based on a four-class prediction model. (***A***) Stacked bar plots indicate the prediction scores of benchmark RNA genes for each of the four classes: dark blue-phenotype protein-coding gene (Ph), cyan-RNA gene (RNA), red-pseudogene (Ps), yellow n random intergenic sequence (Ig). Ig were included to provide another set of likely nonfunctional sequences distinct from pseudogenes. Expression breadth and tissue-specific features were excluded and 100 bp sequences were used. A benchmark RNA gene is classified as one of the four classes according to the highest prediction score. The color bars below the chart indicate the predicted class, with the same color scheme as the prediction score. Sequences classified as Ph or RNA were considered functional, while those classified as Ps or Ig were considered nonfunctional. Percentages below a classification region indicate the proportion of sequences classified as that class. (***B***) Phenotype protein-coding gene prediction scores. (***C***) Pseudogene prediction scores. (***D***) Random unexpressed intergenic region prediction scores. Note that no sequence was predicted as functional. (***E***) Intergenic transcribed region (ITR), (***F***) Araport11 ncRNA regions. (***G***) TAIR10 ncRNA regions. Note that the 100 bp model used here allowed us to evaluate an additional 10,938 ITRs and 1,406 annotated ncRNAs compared to the 500 bp full and tissue-agnostic models.

To determine whether the suboptimal predictions by the phenotype protein-coding gene-based models are because RNA genes belong to a class of their own, we next built a multi-class function prediction model (as opposed to the binary, two-class models described above) aimed at distinguishing four classes of sequences: benchmark RNA genes (n=46, Supplementary Table 6), phenotype protein-coding genes (1,882), pseudogenes (3,916), and randomly-selected, unexpressed intergenic regions (4,000). In the four-class model, 87% of benchmark RNA genes, including all six phenotype RNA genes, were predicted as functional sequences (65% RNA gene-like and 22% phenotype protein-coding gene-like; **Fig. 6A**). In addition, 95% of phenotype protein-coding genes were predicted as functional (**Fig. 6B**), including 80% of narrowly expressed genes, an increase of 22% over the 500 bp tissue-agnostic model (Supplementary Fig. 3C). For benchmark non-functional sequences, 70% of pseudogenes (**Fig. 6C**) and 100% of unexpressed intergenic regions (**Fig. 6D**) were predicted as nonfunctional (either as pseudogenes or unexpressed intergenic sequences). Overall, the four-class model improves prediction accuracy of RNA genes and narrowly expressed genes. In addition, given the 30% FPR among pseudogenes, the four-class model provides a liberal estimate of sequence functionality and high confidence estimate of non-functionality.

**Supplementary Figure 6.**
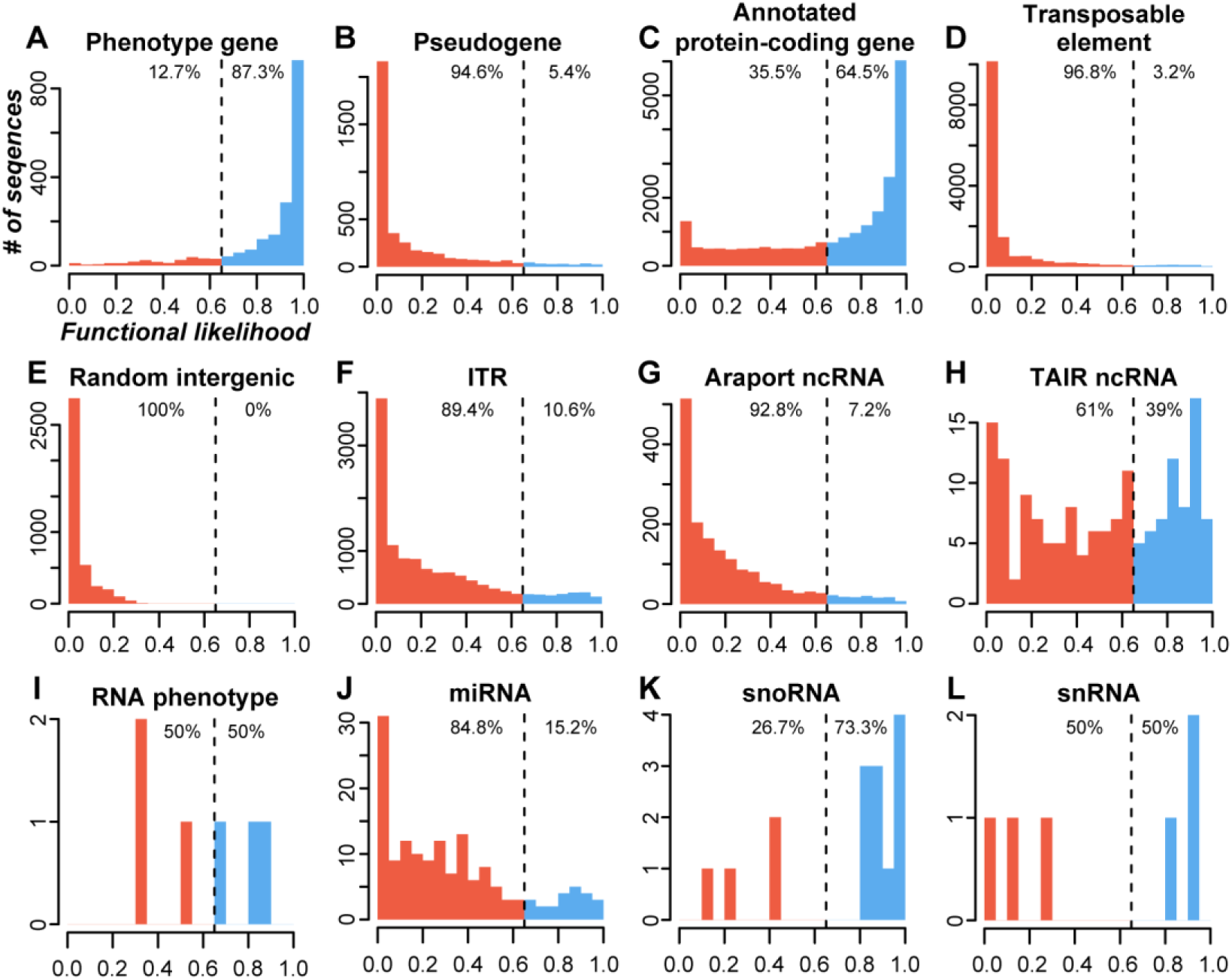
Distributions of functional likelihood scores based on the 100 bp tissue-agnostic model. (*A*) Phenotype genes. (*B*) Pseudogenes. (*C*) Protein-coding gene. (*D*) transposable elements. (*E*) Random unexpressed intergenic sequences. (*F*) Intergenic transcribed regions (ITR). (*G*) Araport11 ncRNAs. (*H*) TAIR10 ncRNAs. (*I*) RNA genes with loss-of-function mutant phenotypes. (J) MicroRNAs, (*K*) Small nucleolar RNAs, (*L*) Small nuclear RNAs. The tissue-agnostic model was built with 100 bp features and while excluding the expression breadth and tissue-specific features as annotated RNA genes tend to be more narrowly expressed than phenotype genes (U tests, all *p* < 2e-05; Supplementary Fig. 2A). Higher functional likelihood values indicate greater similarity to phenotype genes while lower values indicate similarity to pseudogenes. Vertical dashed lines display the threshold to define a sequence as functional or nonfunctional. The numbers to the left and right of the dashed line show the percentage of sequences predicted as functional or nonfunctional, respectively.

### Most intergenic transcribed regions and annotated ncRNAs do not resemble benchmark RNA genes

By applying the four-class model on ITRs and annotated ncRNAs, we found that 34% of ITRs, 38% of Araport ncRNAs, and of 65% TAIR ncRNAs were predicted as functional sequences (**Fig. 6E-G**). Specifically, ≤20% of ITR and annotated ncRNA sequences were classified as RNA genes (**Fig. 6E-G**). Although miRNAs dominate the benchmark RNA sequences, we should emphasize that the four-class prediction model also increased the proportion of snoRNAs and snRNAs that were predicted as functional (91%) compared to the 100bp tissue agnostic model (67%). Thus a lack of similarity to benchmark miRNAs provides evidence that most ITRs and Araport ncRNAs are not functioning as RNA genes.

To provide an overall estimate of the proportion of likely functional and nonfunctional ITRs and annotated ncRNAs, we considered the predictions from the four-class model (**Fig. 6**), the full model (**Fig. 3,4**), and the tissue-agnostic models (Supplementary Fig. 4, 6), which cover both protein-coding and RNA gene functions. Based on support from ≥1 of the four models, we classified 4,437 ITRs (38%) and 796 annotated ncRNAs (44%) as likely functional, as they resembled either phenotype protein-coding or RNA genes. Although our findings lend support that they are likely parts of novel or annotated genes, we should stress that, given the relatively high FPR in the tissue-agnostic and four-class models, the estimate of functional ITR/ncRNA is a liberal one. Most importantly, we find that a substantial number of ITRs (62%) and annotated ncRNAs (56%) are predicted as nonfunctional. Moreover, at least a third of ITRs (**Fig. 6E**) and Araport ncRNAs (**Fig. 6F**) most closely resemble unexpressed intergenic regions. Based on these findings, we conclude that the majority of ITRs and annotated ncRNA regions resemble nonfunctional genomic regions, and therefore are derived from noisy transcription.

## CONCLUSION

Discerning the location of functional regions within a genome represents a key goal in genomic biology and is fundamental to molecular evolutionary studies. Despite advances in computational gene finding, it remains challenging to determine whether ITRs represent functional or noisy biochemical activity. We established robust function prediction models based on the evolutionary, biochemical, and structural characteristics of phenotype genes and pseudogenes in *A. thaliana*. The prediction models accurately define functional and nonfunctional regions and are applicable genome-wide and echo recent findings using human data to evaluate RNA gene functionality (Tsai et al. 2017). We utilized prediction models to assess the functionality of both protein-coding and annotated RNA genes. As benchmark examples of more recently identified RNA gene classes become available in *A. thaliana*, such as *cis*-acting regulatory (Guil and Esteller 2012) or competitive endogenous (Tan et al. 2015) RNAs, it will be interesting to see if sequences that encode RNA products with these roles can be predicted as functional based on a similar predictive framework. Given that function predictions were successful in both plants and metazoans, integrating the evolutionary and biochemical features of known genes for functional genomic region prediction will likely be applicable to any species. The next step will be to test whether function prediction models can be applied across species, which could ultimately allow the phenotype data and omics resources available in model systems to effectively guide the identification of functional regions in non-models.

Expression data was highly informative to functional predictions. We found that the prediction model based on only 24 transcription activity-related features performs nearly as well as the full model that integrates additional information including conservation, H3 mark, methylation, and TF binding data. In human, use of transcription data from cell lines also produced highly accurate predictions of functional genomic regions (Tsai et al. 2017). Importantly, our findings suggest that function prediction models can be established in any species, model or not, with a modest number of transcriptome datasets (e.g. 51 in this study and 19 in human). Despite the importance of transcription data, we emphasize that the presence of expression evidence is an extremely poor predictor of functionality. With the effectiveness of our model noted, one major caveat of our models is that narrowly-expressed phenotype genes are frequently predicted as pseudogene and broadly-expressed pseudogenes tend to be called functional. To improve the function prediction model, it will be important to explore additional features unrelated to transcription, particularly those relevant to broadly expressed pseudogenes that are mostly likely recently pseudogenized. In addition, because few phenotype genes are narrowly-expressed (5%) in the *A. thaliana* training data, more phenotyping data for narrowly expressed genes will be crucial as well.

Upon application of the function prediction models genome-wide, 4,427 ITRs and 796 annotated ncRNAs in *A. thaliana* are predicted as functional sequences. However, considering the high false positive rates (e.g. 10% for the full and 31% for the four-class model), this is most likely an overestimate of the functional sequences contributed by ITRs and annotated ncRNAs. While we err on the side of calling non-functional sequences as functional, we reduce the error rate for calling a functional sequence as non-functional. Despite this conservative approach to classifying sequences as non-functional, the majority of ITRs and ncRNAs resemble pseudogenes and random unexpressed intergenic regions. Similarly, most human ncRNAs are more similar to nonfunctional sequences than they are to protein coding and RNA genes (Tsai et al. 2017). Together with our finding of a significant relationship between the amount of intergenic expression and genome size, we conclude that a significant proportion of intergenic transcripts are nonfunctional noise. Thus, instead of assuming any expressed sequence must be functionally significant, we advocate that the null hypothesis should be that it is not, particularly considering that most ITRs and annotated ncRNAs have not been experimentally characterized.

## MATERIALS AND METHODS

### Identification of transcribed regions in leaf tissue of 15 flowering plants

RNA-sequencing (RNA-seq) datasets were retrieved from the Sequence Read Archive (SRA) at the National Center for Biotechnology Information (NCBI; www.ncbi.nlm.nih.gov/sra/) for 15 flowering plant species (Supplementary Table 1). All datasets were generated from leaf tissue and sequenced on Illumina HiSeq 2000 or 2500 platforms. Genome sequences and gene annotation files were downloaded from Phytozome v.11 (www.phytozome.net) (Goodstein et al. 2012) or Oropetium Base v.01 (www.sviridis.org) (VanBuren et al. 2015). Genome sequences were repeat masked using RepeatMasker v4.0.5 (www.repeatmasker.org) if a repeat-masked version was not available. Only one end from paired-end read datasets were utilized in downstream processing. Reads were trimmed to be rid of low scoring ends and residual adaptor sequences using Trimmomatic v0.33 (LEADING:3 TRAILING:3 SLIDINGWINDOW:4:20 MINLEN:20) (Bolger et al. 2014) and mapped to genome sequences using TopHat v2.0.13 (default parameters except as noted below) (Kim et al. 2013). Reads ≥20 nucleotides in length that mapped uniquely within a genome were used in further analysis.

For each species, thirty million mapped reads were randomly selected from among all datasets and assembled into transcript fragments using Cufflinks v2.2.1 (default parameters except as noted below) (Trapnell et al. 2010), while correcting for sequence-specific biases during the sequencing process by providing an associated genome sequence with the -b flag. The expected mean fragment length for assembled transcript fragments in Cufflinks was set to 150 from the default of 200 so that expression levels in short fragments would not be overestimated. The 1^st^ and 99^th^ percentile of intron lengths for each species were used as the minimum and maximum intron lengths, respectively, for both the TopHat2 and Cufflinks steps. Intergenic transcribed regions (ITRs) were defined by transcript fragments that did not overlap with gene annotation and did not have significant six-frame translated similarity to plant protein sequences in Phytozome v.10 (BLASTX E-value < 1E-05). The correlation between assembled genome size and gene counts was determined with data from the first 50 published plant genomes (Michael and Jackson 2013).

### Phenotype data sources

Mutant phenotype data for *A. thaliana* protein-coding genes was collected from a published dataset (Lloyd and Meinke 2012), the Chloroplast 2010 database (Ajjawi et al. 2010; Savage et al. 2013), and the RIKEN phenome database (Kuromori et al. 2006) as described by Lloyd et al. (2015). Phenotype genes used in our analyses were those whose disruption resulted in lethal or visible defects under standard laboratory growth conditions. Genes with documented mutant phenotypes under standard conditions were considered as a distinct and non-overlapping category from other annotated protein-coding genes. We identified six RNA genes with documented loss-of-function phenotypes through literature searches (Supplementary Table 7): *At4* (AT5G03545) (Shin et al. 2006), *MIR164A* and *MIR164D* (AT2G47585 and AT5G01747, respectively) (Guo et al. 2005), *MIR168A* (AT4G19395) (W. Li et al. 2012), and *MIR828A* and *TAS4* (AT4G27765 and AT3G25795, respectively) (Hsieh et al. 2009). Conditional phenotype genes were those belonging to the Conditional phenotype group as described by Lloyd and Meinke (2012). Loss-of-function mutants of these genes exhibited phenotype only under stress conditions.

### Arabidopsis thaliana genome annotation

*A. thaliana* protein-coding gene, miRNA gene, snoRNA gene, snRNA gene, ncRNA region, pseudogene, and transposable element annotations were retrieved from The Arabidopsis Information Resource v.10 (TAIR10; www.arabidopsis.org) (Berardini et al. 2015). Additional miRNA gene and lncRNA region annotations were retrieved from Araport v.11 (www.araport.org). A primary difference between the TAIR ncRNAs and Araport lncRNAs (referred to as Araport ncRNAs in the Results & Discussion section) is the date in which they were annotated. For example, 221 ncRNAs were present in the v.7 release of TAIR, which dates back to 2007 (TAIR10 contains 394 ncRNA annotations) (Swarbreck et al. 2008; Lamesch et al. 2012; Berardini et al. 2015). However, Araport lncRNAs were annotated in the past five years (Krishnakumar et al. 2015). Thus, that TAIR ncRNAs are generally more highly and broadly expressed is likely a result of the less sensitive transcript identification methods available for early TAIR releases. A pseudogene-finding pipeline (Zou et al. 2009) was used to identify additional pseudogene fragments and count the number of disabling mutations (premature stop or frameshift mutations). Genes, pseudogenes, and transposons with overlapping annotation were excluded from further analysis. Overlapping lncRNA annotations were merged for further analysis. When pseudogenes from TAIR10 and the pseudogene-finding pipeline overlapped, the longer pseudogene annotation was used.

*A. thaliana* ITRs analyzed include: (1) the Set 2 ITRs in Moghe et al. (Moghe et al. 2013), (2) the novel transcribed regions from Araport v.11, and (3) additional ITRs from 206 RNA-seq datasets (Supplementary Table 5). Reads were trimmed, mapped, and assembled into transcript fragments as described above, except that overlapping transcript fragments from across datasets were merged. ITRs analyzed did not overlap with any TAIR10, Araport11, or pseudogene annotation. Overlapping ITRs from different annotated subsets were kept based on a priority system: Araport11 > Set 2 ITRs from Moghe et al. (Moghe et al. 2013) > ITRs identified in this study. For each sequence entry (gene, ncRNA, pseudogene, transposable element, or ITR), a 100 and 500 base pair (bp) window was randomly chosen for calculating feature values and subsequent model building steps. Feature descriptions are provided in the following sections. The feature values for randomly selected 500 and 100 bp windows are provided in Supplementary Tables 2 and 6, respectively. Additionally, non-expressed intergenic sequences were randomly-sampled from genome regions that did not overlap with annotated genes, pseudogenes, transposable elements, or regions with genic or intergenic transcript fragments (100 bp, n=4,000; 500 bp, n=3,716). All 100 and 500 bp windows described above are referred to as sequence windows throughout the Methods section.

### Sequence conservation and structure features

There were 10 sequence conservation features examined. The first two were derived from comparisons between *A. thaliana* accessions including nucleotide diversity and Tajima’s D among 81 accessions (Cao et al. 2011) using a genome matrix file from the 1,001 genomes database (www.1001genomes.org). The python scripts are available through GitHub (https://github.com/ShiuLab/GenomeMatrixProcessing). The remaining eight features were derived from cross-species comparisons, three based on multiple sequence and five based on pairwise alignments. Three multiple sequence alignment-based features were established using aligned genomic regions between *A. thaliana* and six other plant species (*Glycine max*, *Medicago truncatula*, *Populus trichocarpa*, *Vitis vinifera*, *Sorghum bicolor*, and *Oryza sativa*) (F. Li et al. 2012), which are referred to as conserved blocks. For each conserved block, the first feature was the proportion of a sequence window that overlapped a conserved block (referred to as coverage), and the two other features were the maximum and average phastCons scores within each sequence window. The phastCons score was determined for each nucleotide within conserved blocks (F. Li et al. 2012). Nucleotides in a sequence window that did not overlap with a conserved block were assigned a phastCons score of 0. For each sequence window, five pairwise alignment-based cross-species conservation features were the percent identities to the most significant BLASTN match (if E-value<1E-05) in each of five taxonomic groups. The five taxonomic groups included the *Brassicaceae* family (n_species_=7), other dicotyledonous plants (22), monocotyledonous plants (7), other embryophytes (3), and green algae (5). If no sequence with significant similarity was present, percent identity was scored as zero.

For sequence-structure features, we used 125 conformational and thermodynamic dinucleotide properties collected from DiProDB database (Friedel et al. 2009). Because the number of dinucleotide properties was high and dependent, we reduced the dimensionality by utilizing principal component (PC) analysis as described previously (Tsai et al. 2015). Sequence-structure values corresponding to the first five PCs were calculated for all dinucleotides in and averaged across the length of a sequence window and used as features when building function prediction models.

### Transcription activity features

We generated four multi-dataset and 20 individual dataset transcription activity features. To identify a set of RNA-seq datasets to calculate multi-dataset features, we focused on the 72 of 206 RNA-seq datasets each with ≥20 million reads (see above; Supplementary Table 5). Transcribed regions were identified with TopHat2 and Cufflinks as described in the RNA-seq analysis section except that the 72 *A. thaliana* RNA-seq datasets were used. Following transcript assembly, we excluded 21 RNA-seq datasets because they had unusually high RPKM (Reads Per Kilobase of transcript per Million mapped reads) values (median RPKM value range=272∼2,504,294) compared to the rest (2∼252). The remaining 51 RNA-seq datasets were used to generate four multi-dataset transcription activity features including: expression breadth, 95^th^ percentile expression level, maximum transcript coverage, and presence of expression evidence (for values see Supplementary Table 2, 6). Expression breadth was the number of RNA-seq datasets that have ≥1 transcribed region that overlapped with a sequence window. The 95^th^ percentile expression level was the 95^th^ percentile of RPKM values across 51 RNA-seq datasets where RPKM values were set to 0 if there was no transcribed region for a sequence window. Maximum transcript coverage was the maximum proportion of a sequence window that overlapped with a transcribed region across 51 RNA-seq datasets. Presence of expression evidence was determined by overlap between a sequence window and any transcribed region in the 51 RNA-seq datasets.

In addition to features based on multiple datasets, 20 individual dataset features were derived from 10 datasets: seven tissue/organ-specific RNA-seq datasets including pollen (SRR847501), seedling (SRR1020621), leaf (SRR953400), root (SRR578947), inflorescence (SRR953399), flower, (SRR505745) and silique (SRR953401), and three datasets from non-standard growth conditions, including dark-grown seedlings (SRR974751) and leaf tissue under drought (SRR921316) and fungal infection (SRR391052). For each of these 10 RNA-seq datasets, we defined two features for each sequence window: the maximum transcript coverage (as described above) and the maximum RPKM value of overlapping transcribed regions (referred to as Level in **Fig. 2**). If no transcribed regions overlapped a sequence window, the maximum RPKM value was set as 0. For the analysis of narrowly- and broadly-expressed phenotype genes and pseudogenes (Supplementary Fig. 3B,C), we used 28 out of 51 RNA-seq datasets generated from a single tissue and in standard growth conditions to calculate the number of tissues with evidence of expression (tissue expression breadth). In total, seven tissues were represented among the 28 selected RNA-seq datasets (see above; Supplementary Table 5), and thus tissue expression breadth ranges from 0 to 7 (note that only 1 through 7 are shown in Supplementary Fig. 3B,C due to low sample size of phenotype genes in the 0 bin). The tissue breadth value is distinct from the expression breadth feature used in model building that was generated using all 51 datasets and considered multiple RNA-seq datasets from the same tissue separately (range: 0-51).

### Histone 3 mark features

Twenty histone 3 (H3) mark features were calculated based on eight H3 chromatin immunoprecipitation sequencing (ChIP-seq) datasets from SRA. The H3 marks examined include four associated with activation (H3K4me1: SRR2001269, H3K4me3: SRR1964977, H3K9ac: SRR1964985, and H3K23ac: SRR1005405) and four associated with repression (H3K9me1: SRR1005422, H3K9me2: SRR493052, H3K27me3: SRR3087685, and H3T3ph: SRR2001289). Reads were trimmed as described in the RNA-seq section and mapped to the TAIR10 genome with Bowtie v2.2.5 (default parameters) (Langmead et al. 2009). Spatial Clustering for Identification of ChIP-Enriched Regions v.1.1 (Xu et al. 2014) was used to identify ChIP-seq peaks with a false discover rate ≤ 0.05 with a non-overlapping window size of 200, a gap parameter of 600, and an effective genome size of 0.92 (Koehler et al. 2011). For each H3 mark, two features were calculated for each sequence window: the maximum intensity among overlapping peaks and peak coverage (proportion of overlap with the peak that overlaps maximally with the sequence window). In addition, four multi-mark features were generated. Two of the multi-mark features were the number of activating marks (0-4) overlapping a sequence window and the proportion of a sequence window overlapping any peak from any of the four activating marks (activating mark peak coverage). The remaining two multi-mark features were the same as the two activating multi-mark features except focused on the four repressive marks.

### DNA methylation features

Twenty-one DNA methylation features were calculated from bisulfite-sequencing (BS-seq) datasets from seven tissues (pollen: SRR516176, embryo: SRR1039895, endosperm: SRR1039896, seedling: SRR520367, leaf: SRR1264996, root: SRR1188584, and inflorescence: SRR2155684). BS-seq reads were trimmed as described above and processed with Bismark v.3 (default parameters) (Krueger and Andrews 2011) to identify methylated and unmethylated cytosines in CG, CHH, and CHG (H = A, C, or T) contexts. Methylated cytosines were defined as those with ≥5 mapped reads and with >50% of mapped reads indicating that the position was methylated. For each BS-seq dataset, the percentage of methylated cytosines in each sequence window for CG, CHG, and CHH contexts were calculated if the sequence window had ≥5 cytosines with ≥5 reads mapping to the position. To determine whether the above parameters where reasonable, we assessed the false positive rate of DNA methylation calls by evaluating the proportion of cytosines in the chloroplast genome that are called as methylated, as the chloroplast genome has few DNA methylation events (Ngernprasirtsiri et al. 1988; Zhang et al. 2006). Based on the above parameters, 0-1.5% of cytosines in CG, CHG, or CHH contexts in the chloroplast genome were considered methylated in any of the seven BS-seq datasets. This indicated that the false positive rates for DNA methylation calls were low and the parameters were reasonable.

### Chromatin accessibility and transcription factor binding features

Chromatin accessibility features consisted of ten DHS-related features and one micrococcal nuclease sequencing (MNase-seq)-derived feature. DHS peaks from five tissues (seed coat, seedling, root, unopened flowers, and opened flowers) were retrieved from the Gene Expression Omnibus (GSE53322 and GSE53324) (Sullivan et al. 2014). For each of the five tissues, the maximum DHS peak intensity and DHS peak coverage were calculated for each sequence window. Normalized nucleosome occupancy per bp based on MNase-seq was obtained from Liu et al. (Liu et al. 2015). The average nucleosome occupancy value was calculated across each sequence window. Transcription factor (TF) binding site features were based on *in vitro* DNA affinity purification sequencing data of 529 TFs (O’Malley et al. 2016). Two features were generated for each sequence window: the total number of TF binding sites and the number of distinct TFs bound.

### Single-feature prediction performance

The ability for each single feature to distinguish between functional and nonfunctional regions was evaluated by calculating AUC-ROC value with the Python scikit-learn package (Pedregosa et al. 2011). Thresholds to predict sequences as functional or nonfunctional using a single feature were defined by the feature value that produced the highest F-measure, the harmonic mean of precision (proportion of sequences predicted as functional that are truly functional) and recall (proportion of truly functional sequences predicted as functional). The F-measure allows consideration of both false positives and false negatives at a given threshold. FPR were calculated as the percentage of negative (nonfunctional) cases with values above or equal to the threshold and thus falsely predicted as functional. FNR were calculated as the percentage of positive (functional) cases with values below the threshold and thus falsely predicted as nonfunctional.

### Binary classification with machine learning

For binary classification (two-class) models that contrasted phenotype genes and pseudogenes, the random forest (RF) implementation in the Waikato Environment for Knowledge Analysis software (WEKA) (Hall et al. 2009) was utilized. Three types of two-class models were established, including the full model (500 bp sequence window, **Fig. 3A,B** and **Fig. 4**), tissue-agnostic models (500 bp, Supplementary Fig. 4; 100 bp, Supplementary Fig. 6), and single feature category models (**Fig. 3A,B**). For each model type, we first generated 100 balanced datasets by randomly selecting equal numbers of phenotype genes (positive examples) and pseudogenes (negative examples). For each of these 100 datasets, 10-fold stratified cross-validation was utilized, where the model was trained using 90% of sequences and tested on the remaining 10%. Thus, for each model type, a sequence window had 100 prediction scores, where each score was the proportion of 500 random forest trees that predicted a sequence as a phenotype gene in a balanced dataset. The median of 100 prediction scores was used as the functional likelihood (FL) value (Supplementary Table 4). The FL threshold to predict a sequence as functional or nonfunctional was defined based on maximum F-measure as described in the previous section.

We tested multiple-K parameters (2 to 25) in the WEKA-RF implementation, which alters the number of randomly-selected features included in each RF tree (Supplementary Table 8), and found that 15 randomly-selected features provided the highest performance based on AUC-ROC (calculated and visualized using the ROCR package) (Sing et al. 2005). Feature importance was assessed by excluding one feature at a time to determine the associated reduction in prediction performance (Supplementary Table 9). All leave-one-out models performed well (AUC-ROC >0.97), indicating that no single feature was dominating the function predictions and/or many features are correlated (Supplementary Fig. 7). Binary classification models were also built using all features from 500 bp sequences (equivalent to the full model) with the Sequential Minimal Optimization - Support Vector Machine (SMO-SVM) implementation in WEKA (Hall et al. 2009). The results of SMO-SVM models were highly similar to the full RF results: *PCC* between the FL values generated by RF and SMO-SVM=0.97; AUC-ROC of SMO-SVM=0.97; FPR=12%; FNR=3%. By comparison, the full RF model had AUC-ROC=0.98, FPR=10%, FNR=4%.

**Supplementary Figure 7.**
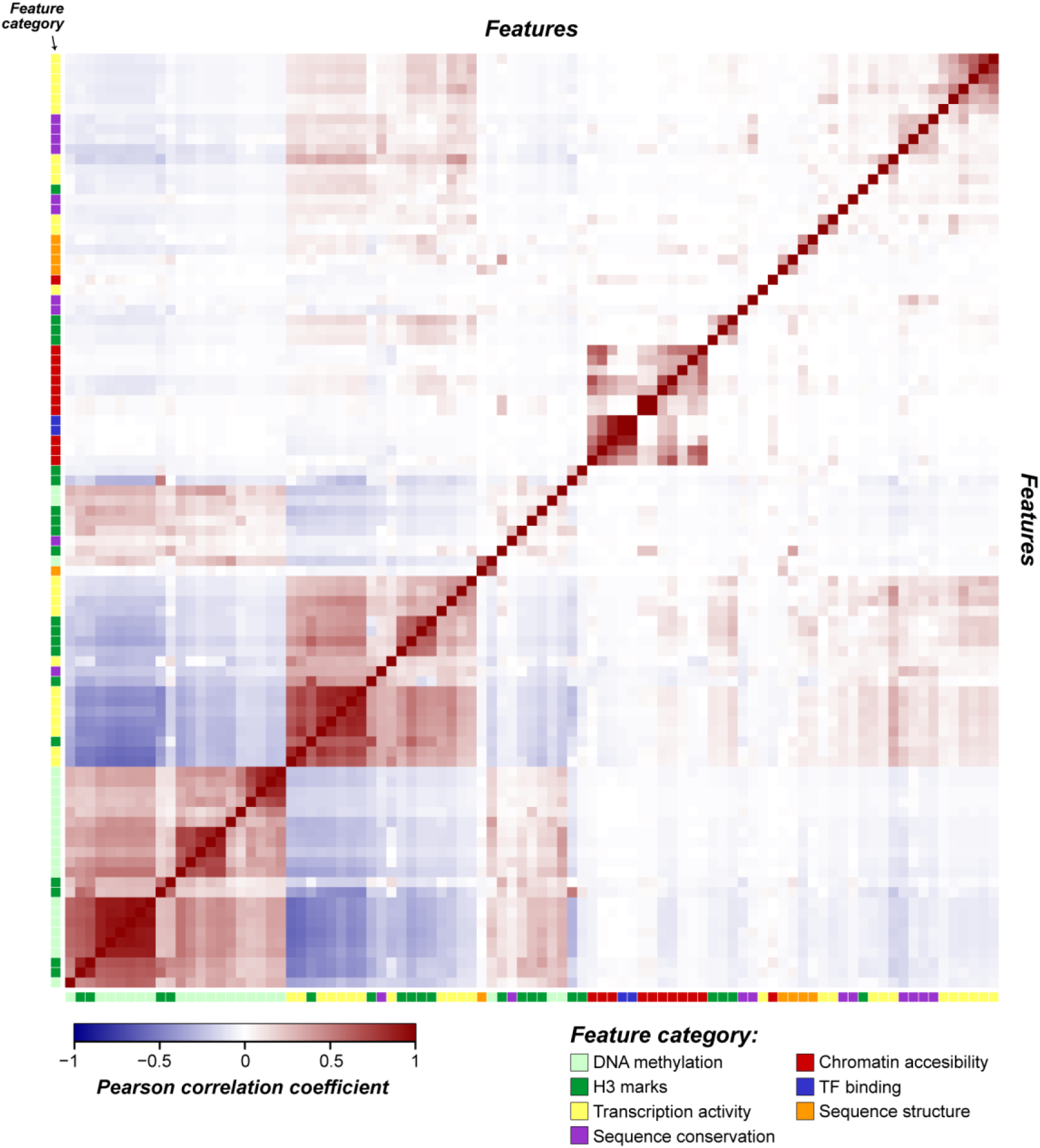
Correlation between features used in functional predictions. Colors within the heatmap indicate pairwise correlation between two features. Colors on the left-most and bottom-most edges indicate the associated feature category (Fig. 2). Feature values were quantile normalized prior to calculating correlation.

Tissue-agnostic models were generated by excluding the expression breadth feature and 95^th^ percentile expression level and replacing all features from RNA-seq, BS-seq, and DHS datasets that were available in multiple tissues. For multiple-tissue RNA-seq data, the maximum expression level across 51 RNA-seq datasets (in RPKM) and maximum coverage (as described in the transcription activity section) of a sequence window in any of 51 RNA-seq datasets were used. For multi-tissue DNA methylation features, minimum proportions of methylated cytosines in any tissue in CG, CHG, and CHH contexts were used. For DHS data, the maximum peak intensity and peak coverage was used instead. In single feature category predictions, fewer total features were used and therefore lower–K values (i.e. the number of random features selected when building random forests) were considered in parameter searches (Supplementary Table 8).

### Multi-class machine learning model

For the four-class model, benchmark RNA gene, phenotype protein-coding gene, pseudogene, and random unexpressed intergenic sequences were used as the four training classes. Benchmark RNA genes consisted of six RNA genes with documented loss-of-function phenotypes and 40 high-confidence miRNA genes from miRBase (www.mirbase.org) (Kozomara and Griffiths-Jones 2014). We considered that the decreased numbers of benchmark RNA genes would not allow us to effectively distinguish between sequence classes. However, binary predictions generated using 35 phenotype gene and pseudogene instances and the 100 bp tissue-agnostic feature set resulted in an AUC-ROC performance of 0.96. We generated 250 datasets with equal proportions (larger classes randomly sampled) of training sequences. Twofold stratified cross-validation was utilized due to the low number of benchmark RNA genes. The features included those described for the tissue-agnostic model and focused on 100 bp sequence windows. The RF implementation, *cforest*, in the *party* package of R (Strobl et al. 2008) was used to build the classifiers. The four-class predictions provide prediction scores for each sequence type: an RNA gene, phenotype protein-coding gene, pseudogene, and unexpressed intergenic score (Supplementary Table 4). The prediction scores indicate the proportion of random forest trees that classify a sequence as a particular class. Median prediction scores from across 100 balanced runs were used as final prediction scores. Scores from a single balanced dataset models sum to 1, but not the median from 100 balanced runs. Thus, the median scores were scaled to sum to 1. For each sequence window, the maximum prediction score among the four classes was used to classify a sequence as phenotype gene, pseudogene, unexpressed intergenic region, or RNA gene.

## AVAILABILITY

All relevant data are within the article and supplementary data files.

## SUPPLEMENTARY MATERIALS

Supplementary Data are available online.

## ACKNOWLEDGEMENTS

The authors wish to thank Christina Azodi, Ming-Jung Liu, Gaurav Moghe, Bethany Moore, and Sahra Uygun and for providing processed data and discussion. This work was supported by the National Science Foundation (grant numbers IOS-1126998, IOS-1546617, and DEB-1655386 to S.H.S), and Research Experience for Undergraduates support [to R.P.S]; and the Michigan State University Dissertation Continuation Fellowship [to J.P.L].

## SUPPLEMENTARY TABLES

**Supplementary Table 1.** Leaf tissue RNA-sequencing datasets for 15 flowering plant species

**Supplementary Table 2.** Conservation, biochemical, and sequence-structure feature values calculated from 500 bp sequences.

**Supplementary Table 3.** False positive and false negative rates for single feature classifications.

**Supplementary Table 4.** Function predictions for all models generated in this study.

**Supplementary Table 5.** RNA-sequencing datasets for identifying intergenic transcribed regions, calculating transcription activity features, and assessing tissue-specific predictions.

**Supplementary Table 6.** Conservation, biochemical, and sequence-structure feature values calculated from 100 bp sequences.

**Supplementary Table 7.** RNA genes with documented loss-of-function phenotypes.

**Supplementary Table 8.** K parameters tested for random forest runs.

**Supplementary Table 9.** AUC-ROC scores from leave-one-feature-out machine learning runs.

## SUPPLEMENTARY INFORMATION

### Variability in single feature performance within feature categories

Within each feature category, there was a wide range of performance between features (**Fig. 2**, Supplementary Table 3) and there were clear biological or technical explanations for features that perform poorly. For the transcription activity category, 17 out of 24 features had an AUC-ROC performance >0.8, including the best-performing feature, expression breadth (AUC-ROC=0.95; **Fig. 2A**). However, five transcription activity-related features performed poorly (AUC-ROC<0.65), including the presence of expression (transcript) evidence (AUC-ROC=0.58; **Fig. 2A**). For the sequence conservation category, maximum and average phastCons conservation scores were highly distinct between phenotype genes and pseudogenes (AUC-ROC=0.83 and 0.82, respectively; **Fig. 2B**). On the other hand, identity to best matching nucleotide sequences found in *Brassicaceae* and algal species were not informative (AUCROC=0.55 and 0.51, respectively; **Fig. 2B**). This was because 99.8% and 95% of phenotype genes and pseudogenes, respectively, had a potentially homologous sequence within the *Brassicaceae* family, and only 3% and 1%, respectively, in algal species. Thus, *Brassicaceae* genomes were too similar and algal genomes too dissimilar to *A. thaliana* to provide meaningful information. H3 mark features also displayed high variability. The most informative H3 mark features were based on the number and coverage of activation-related marks (AUC-ROC=0.87 and 0.85, respectively; **Fig. 2E**), consistent with the notion that histone marks are often jointly associated with active genomic sequences to provide a robust regulatory signal (Schreiber and Bernstein 2002; Wang et al. 2008). By comparison, the coverage and intensity of H3 lysine 27 trimethylation (H3K27me3) and H3 threonine 3 phosphorylation (H3T3ph) were largely indistinct between phenotype genes and pseudogenes (AUC-ROC range: 0.55-0.59; **Fig. 2E**).

### Error rates from single-feature functional predictions

The differences between genes and pseudogenes in transcription, conservation, and epigenetic features and functional genomic regions suggested that these features may individually provide sufficient information for distinguishing between functional and nonfunctional genomic regions. To assess this possibility, we next evaluated the error rates of function predictions based on single features. We first considered expression breadth of a sequence, the best predicting single feature of functionality. Despite high AUC-ROC (0.95; **Fig. 2A**), the false positive rate (FPR; % of pseudogenes predicted as phenotype genes) was 21% when only expression breadth was used, while the false negative rate (FNR; % of phenotype genes predicted as pseudogenes) was 4%. Similarly, the best-performing H3 mark- and sequence conservation-related features (**Fig.2B,E**) had FPRs of 26% and 32%, respectively, and also incorrectly classified at least 10% of phenotype genes as pseudogenes. Thus, error rates are high even when considering well-performing single features, indicating the need to jointly consider multiple features for distinguishing phenotype genes and pseudogenes.

### Features of misclassified sequences

Although the full model performs exceedingly well, there remain false predictions. There are 76 phenotype genes (4%) predicted as nonfunctional (referred to as low-functional likelihood (FL) phenotype genes). We assessed why these phenotype genes were not correctly identified by first asking what category of features were particularly distinct between low-FL and the remaining phenotype genes. We found that the major category that led to the misclassification of phenotype genes was transcription activity, as only 7% of low-scoring phenotype genes were predicted as functional in the transcription activity-only model, compared to 98% of high FL phenotype genes (**Fig. 5**). By contrast, >65% of low-FL phenotype genes were predicted as functional when sequence conservation, H3 mark, or DNA methylation features were used. This could suggest that the full model is less effective in predicting functional sequences that are weakly or narrowly expressed. While sequence conservation features are distinct between functional and nonfunctional sequences when considered in combination, a significantly higher proportion of low-FL phenotype genes were specific to the *Brassicaceae* family, with only 33% present in dicotyledonous species outside of the *Brassicaceae*, compared to 78% of high-scoring phenotype genes (FET, *p* < 4e-12), thus our model likely has reduced power in detecting lineage-specific sequences.

We also predict 80 pseudogenes (10%) to be functional (high-FL pseudogenes). A significantly higher proportion of high-FL pseudogenes came from existing genome annotation as 19% of annotated pseudogenes were classified as functional, compared to 4% of pseudogenes identified through a computational pipeline (FET, *p* < 1.5E-10) (Zou et al. 2009). We found that high-FL pseudogenes might be more recently pseudogenized and thus have not yet lost many genic signatures, as the mean number of disabling mutations (premature stop or frameshift) per kb in high-scoring pseudogenes (1.9) were significantly lower than that of low-scoring pseudogenes (4.0; U test, *p* < 0.02). Lastly, we cannot rule out the possibility that a small subset of high-scoring pseudogenes represent truly functional sequences, rather than false positives (Poliseno et al. 2010; Karreth et al. 2015). Overall, the misclassification of both narrowly-expressed phenotype genes and broadly-expressed pseudogenes highlights the need for an updated prediction model that is less influenced by expression breadth.

Among protein-coding genes without phenotype information, we predict 20% as nonfunctional. We expect that at least 4% represent false negatives based on the FNR of the full model. The actual FNR among protein-coding genes may be higher, however, as phenotype genes represent a highly active and well conserved subset of all genes. However, a subset of the low-scoring protein-coding genes may also represent gene sequences undergoing functional decay and *en route* to pseudogene status. To assess this possibility, we examined 1,940 *A. thaliana “*decaying” genes that may be experiencing pseudogenization due to promoter disablement (Yang et al. 2011) and found that, while these decaying genes represented only 7% of all *A. thaliana* annotated protein-coding genes, they made up 45% of protein-coding genes predicted as nonfunctional (Fisher’s Exact Test (FET), *p* < 1E-11).

### Feature correlation between likely-functional ITRs and ncRNAs and their neighboring genes

ITRs and annotated ncRNAs closer to annotated genes tended to be predicted as functional (Supplementary Fig. 5A), as 57% of likely functional and 35% of likely nonfunctional ITRs and ncRNAs were proximal to neighboring genes (within the 95^th^ percentile of intron lengths for all genes) (FET, *p* < 2E-09). Likely functional ITRs and annotated ncRNAs that are proximal to genes may frequently represent unannotated exon extensions. If this is the case, it may be expected that such sequences would exhibit similar features as gene neighbors. However, no clear pattern of increased feature correlation was observed between likely functional ITRs / ncRNAs and neighboring genes when compared to likely non-functional ITRs / ncRNAs or random sequence pairs, regardless of proximity (Supplementary Fig. 5B-D). Thus, despite their proximity to annotated genes, it remains unclear if some ITRs or annotated ncRNAs represent unannotated exon extensions of known genes or not. In addition, for proximal functional ITRs/annotated ncRNAs, we cannot rule out the possibility that they represent false-positive functional predictions due to the accessible and active chromatin states of nearby genes. Given the challenge in ascertaining the origin of likely functional, proximal ITRs/ncRNAs, we instead conservatively estimate that 187 distal, functional ITRs and annotated ncRNAs may represent fragments of novel genes.

## REFERENCES

Ajjawi I, Lu Y, Savage LJ, Bell SM, Last RL. 2010. Large-scale reverse genetics in *Arabidopsis*: case studies from the Chloroplast 2010 Project. Plant Physiol. 152:529–540.

Amundson R, Lauder G V. 1994. Function without purpose. Biol. Philos. 9:443–469.

Berardini TZ, Reiser L, Li D, Mezheritsky Y, Muller R, Strait E, Huala E. 2015. The Arabidopsis information resource: Making and mining the gold standard annotated reference plant genome. Genesis 53:474–485.

Bernard D, Prasanth K V, Tripathi V, Colasse S, Nakamura T, Xuan Z, Zhang MQ, Sedel F, Jourdren L, Coulpier F, et al. 2010. A long nuclear-retained non-coding RNA regulates synaptogenesis by modulating gene expression. EMBO J. 29:3082–3093.

Boeck ME, Huynh C, Gevirtzman L, Thompson OA, Wang G, Kasper DM, Reinke V, Hillier LW, Waterston RH. 2016. The time-resolved transcriptome of C. elegans. Genome Res. 26:1441–1450.

Bolger AM, Lohse M, Usadel B. 2014. Trimmomatic: a flexible trimmer for Illumina sequence data. Bioinformatics 30:2114–2120.

Brown JB, Boley N, Eisman R, May GE, Stoiber MH, Duff MO, Booth BW, Wen J, Park S, Suzuki AM, et al. 2014. Diversity and dynamics of the Drosophila transcriptome. Nature 512:393–399.

Cao J, Schneeberger K, Ossowski S, Günther T, Bender S, Fitz J, Koenig D, Lanz C, Stegle O, Lippert C, et al. 2011. Whole-genome sequencing of multiple Arabidopsis thaliana populations. Nat. Genet. 43:956–963.

Doolittle WF, Brunet TDP, Linquist S, Gregory TR. 2014. Distinguishing between function and effect in genome biology. Genome Biol. Evol. 6:1234–1237.

Eddy SR. 2013. The ENCODE project: missteps overshadowing a success. Curr. Biol. 23:R259–61.

ENCODE Project Consortium. 2012. An integrated encyclopedia of DNA elements in the huma genome. Nature 489:57–74.

Fei Q, Xia R, Meyers BC. 2013. Phased, Secondary, Small Interfering RNAs in Posttranscriptional Regulatory Networks. Plant Cell 25:2400–2415.

Friedel M, Nikolajewa S, Sühnel J, Wilhelm T. 2009. DiProDB: a database for dinucleotide properties. Nucleic Acids Res. 37:D37–-40.

Goodstein DM, Shu S, Howson R, Neupane R, Hayes RD, Fazo J, Mitros T, Dirks W, HellstenU, Putnam N, et al. 2012. Phytozome: a comparative platform for green plant genomics. Nucleic Acids Res. 40:D1178 &86.

Graur D, Zheng Y, Price N, Azevedo RBR, Zufall R a, Elhaik E. 2013. On the immortality of television sets: function in the human genome according to the evolution-free gospel of ENCODE. Genome Biol. Evol. 5:578–590.

Guil S, Esteller M. 2012. Cis-acting noncoding RNAs: friends and foes. Nat. Struct. Mol. Biol. 19:1068–1075.

Gulko B, Gronau I, Hubisz MJ, Siepel A. 2014. Probabilities of Fitness Consequences for Point Mutations Across the Human Genome.

Guo H -S, Xie Q, Fei J -F, Chua N-H. 2005. MicroRNA directs mRNA cleavage of the transcription factor NAC1 to downregulate auxin signals for arabidopsis lateral root development. Plant Cell 17:1376–1386.

Hall M, Frank E, Holmes G, Pfahringer B, Reutemann P, Witten IH. 2009. The WEKA data mining software. ACM SIGKDD Explor. Newsl. 11:10.

Hardiman KE, Brewster R, Khan SM, Deo M, Bodmer R. 2002. The bereft gene, a potential target of the neural selector gene cut, contributes to bristle morphogenesis. Genetics 161:231–247.

Hsieh L -C, Lin S -I, Shih AC -C, Chen J -W, Lin W -Y, Tseng C -Y, Li W -H, Chiou T-J. 2009. Uncovering small RNA-mediated responses to phosphate deficiency in Arabidopsis by deep sequencing. Plant Physiol. 151:2120–2132.

Karreth FA, Reschke M, Ruocco A, Ng C, Chapuy B, Léopold V, Sjoberg M, Keane TM, Verma A, Ala U, et al. 2015. The BRAF pseudogene functions as a competitive endogenous RNA and induces lymphoma in vivo. Cell 161:319–332.

Kellis M, Wold B, Snyder MP, Bernstein BE, Kundaje A, Marinov GK, Ward LD, Birney E, Crawford GE, Dekker J, et al. 2014. Defining functional DNA elements in the human genome. Proc. Natl. Acad. Sci. USA 111:6131–6138.

Kim D, Pertea G, Trapnell C, Pimentel H, Kelley R, Salzberg SL. 2013. TopHat2: accurate alignment of transcriptomes in the presence of insertions, deletions and gene fusions. Genome Biol. 14:R36.

Koehler R, Issac H, Cloonan N, Grimmond SM. 2011. The uniqueome: a mappability resource for short-tag sequencing. Bioinformatics 27:272–274.

Kozomara A, Griffiths -Jones S. 2014. miRBase: annotating high confidence microRNAs using deep sequencing data. Nucleic Acids Res. 42:D68–-73.

Krishnakumar V, Hanlon MR, Contrino S, Ferlanti ES, Karamycheva S, Kim M, Rosen BD, Cheng C -Y, Moreira W, Mock SA, et al. 2015. Araport: the Arabidopsis information portal. Nucleic Acids Res. 43:D1003–-9.

Krueger F, Andrews SR. 2011. Bismark: a flexible aligner and methylation caller for BisulfiteSeq applications. Bioinformatics 27:1571–1572.

Kuromori T, Wada T, Kamiya A, Yuguchi M, Yokouchi T, Imura Y, Takabe H, Sakurai T, Akiyama K, Hirayama T, et al. 2006. A trial of phenome analysis using 4000 *Ds* -insertional mutants in gene-coding regions of Arabidopsis. Plant J. 47:640–651.

Lai K -MV, Gong G, Atanasio A, Rojas J, Quispe J, Posca J, White D, Huang M, Fedorova D, Grant C, et al. 2015. Diverse Phenotypes and Specific Transcription Patterns in Twenty Mouse Lines with Ablated LincRNAs. PLoS One 10:e0125522.

Lamesch P, Berardini TZ, Li D, Swarbreck D, Wilks C, Sasidharan R, Muller R, Dreher K, Alexander DL, Garcia -Hernandez M, et al. 2012. The Arabidopsis Information Resource (TAIR): improved gene annotation and new tools. Nucleic Acids Res. 40:D1202 &10.

Langmead B, Trapnell C, Pop M, Salzberg SL. 2009. Ultrafast and memory-efficient alignment of short DNA sequences to the human genome. Genome Biol. 10:R25.

Li F, Zheng Q, Vandivier LE, Willmann MR, Chen Y, Gregory BD. 2012. Regulatory impact of RNA secondary structure across the Arabidopsis transcriptome. Plant Cell 24:4346–4359.

Li W, Cui X, Meng Z, Huang X, Xie Q, Wu H, Jin H, Zhang D, Liang W. 2012. Transcriptional regulation of Arabidopsis MIR168a and argonaute1 homeostasis in abscisic acid and abiotic stress responses. Plant Physiol. 158:1279–1292.

Li W, Gojobori T, Nei M. 1981. Pseudogenes as a paradigm of neutral evolution. Nature.

Liu M-J, Seddon AE, Tsai ZT -Y, Major IT, Floer M, Howe GA, Shiu S-H. 2015. Determinants of nucleosome positioning and their influence on plant gene expression. Genome Res. 25:1182–1195.

Lloyd J, Meinke D. 2012. A comprehensive dataset of genes with a loss-of-function mutant phenotype in *Arabidopsis*. Plant Physiol. 158:1115–1129.

Lloyd JP, Seddon AE, Moghe GD, Simenc MC, Shiu S-H. 2015. Characteristics of Plant Essential Genes Allow for within- and between-Species Prediction of Lethal Mutant Phenotypes. Plant Cell 27:2133–2147.

Marahrens Y, Panning B, Dausman J, Strauss W, Jaenisch R. 1997. Xist-deficient mice are defective in dosage compensation but not spermatogenesis. Genes Dev. 11:156–166.

Michael TP, Jackson S. 2013. The First 50 Plant Genomes. Plant Genome 6:0.

Moghe GD, Lehti-Shiu MD, Seddon AE, Yin S, Chen Y, Juntawong P, Brandizzi F, Bailey -Serres J, Shiu S-H. 2013. Characteristics and significance of intergenic polyadenylated RNA transcription in *Arabidopsis*. Plant Physiol. 161:210–224.

Nagalakshmi U, Wang Z, Waern K, Shou C, Raha D, Gerstein M, Snyder M. 2008. The Transcriptional Landscape of the Yeast Genome Defined by RNA Sequencing. Science 320:1344–1349.

Neander K. 1991. Functions as selected effects: The conceptual analyst’s defense. Philos. Sci. 58:168–184.

Ngernprasirtsiri J, Kobayashi H, Akazawa T. 1988. DNA methylation as a mechanism of transcriptional regulation in nonphotosynthetic plastids in plant cells. Proc. Natl. Acad. Sci. U. S. A. 85:4750–4754.

Ning S, Wang P, Ye J, Li X, Li R, Zhao Z, Huo X, Wang L, Li F, Li X. 2013. A global map for dissecting phenotypic variants in human lincRNAs. Eur. J. Hum. Genet. 21:1128–1133.

Niu D -K, Jiang L. 2013. Can ENCODE tell us how much junk DNA we carry in our genome? Biochem. Biophys. Res. Commun. 430:1340–1343.

Nobuta K, Venu RC, Lu C, Beló A, Vemaraju K, Kulkarni K, Wang W, Pillay M, Green PJ, Wang G -L, et al. 2007. An expression atlas of rice mRNAs and small RNAs. Nat. Biotechnol. 25:473–477.

O’Malley RC, Huang S -SC, Song L, Lewsey MG, Bartlett A, Nery JR, Galli M, Gallavotti A, Ecker JR. 2016. Cistrome and Epicistrome Features Shape the Regulatory DNA Landscape. Cell 166:1598.

Pedregosa F, Varoquaux G, Gramfort A, Michel V, Thirion B, Grisel O, Blondel M, Prettenhofer P, Weiss R, Dubourg V, et al. 2011. Scikit-learn: Machine Learning in Python. J. Mach. Learn. Res. 12:2825–2830.

Penny GD, Kay GF, Sheardown SA, Rastan S, Brockdorff N. 1996. Requirement for Xist in X chromosome inactivation. Nature 379:131–137.

Poliseno L, Salmena L, Zhang J, Carver B, Haveman WJ, Pandolfi PP. 2010. A coding-independent function of gene and pseudogene mRNAs regulates tumour biology. Nature 465:1033–1038.

Ponting CP, Belgard TG. 2010. Transcribed dark matter: meaning or myth? Hum. Mol. Genet. 19:R162 &8.

Sauvageau M, Goff LA, Lodato S, Bonev B, Groff AF, Gerhardinger C, Sanchez-Gomez DB, Hacisuleyman E, Li E, Spence M, et al. 2013. Multiple knockout mouse models reveal lincRNAs are required for life and brain development. Elife 2:e01749.

Savage LJ, Imre KM, Hall DA, Last RL. 2013. Analysis of essential Arabidopsis nuclear genes encoding plastid-targeted proteins. PLoS One 8:e73291.

Schreiber SL, Bernstein BE. 2002. Signaling Network Model of Chromatin. Cell 111:771–778.

Shin H, Shin H -S, Chen R, Harrison MJ. 2006. Loss of At4 function impacts phosphate distribution between the roots and the shoots during phosphate starvation. Plant J. 45:712n 726.

Simon SA, Meyers BC. 2011. Small RNA-mediated epigenetic modifications in plants. Curr. Opin. Plant Biol. 14:148–155.

Sing T, Sander O, Beerenwinkel N, Lengauer T. 2005. ROCR: visualizing classifier performance in R. Bioinformatics 21:3940–3941.

Stolc V, Samanta MP, Tongprasit W, Sethi H, Liang S, Nelson DC, Hegeman A, Nelson C, Rancour D, Bednarek S, et al. 2005. Identification of transcribed sequences in Arabidopsis thaliana by using high-resolution genome tiling arrays. Proc. Natl. Acad. Sci. U. S. A. 102:4453–4458.

Strobl C, Boulesteix A -L, Kneib T, Augustin T, Zeileis A. 2008. Conditional variable importance for random forests. BMC Bioinformatics 9:307.

Struhl K. 2007. Transcriptional noise and the fidelity of initiation by RNA polymerase II. Nat. Struct. Mol. Biol. 14:103–105.

Sullivan AM, Arsovski AA, Lempe J, Bubb KL, Weirauch MT, Sabo PJ, Sandstrom R, Thurman RE, Neph S, Reynolds AP, et al. 2014. Mapping and dynamics of regulatory DNA and transcription factor networks in A. thaliana. Cell Rep. 8:2015–2030.

Svensson O, Arvestad L, Lagergren J. 2006. Genome-wide survey for biologically functional pseudogenes. PLoS Comput. Biol. 2:e46.

Swarbreck D, Wilks C, Lamesch P, Berardini TZ, Garcia -hernandez M, Foerster H, Li D, Meyer T, Muller R, Ploetz L, et al. 2008. The Arabidopsis Information Resource (TAIR): gene structure and function annotation. Nucleic Acids Res. 36:1009–1014.

Tan JY, Sirey T, Honti F, Graham B, Piovesan A, Merkenschlager M, Webber C, Ponting CP, Marques AC. 2015. Extensive microRNA-mediated crosstalk between lncRNAs and mRNAs in mouse embryonic stem cells. Genome Res. 25:655–666.

Trapnell C, Williams BA, Pertea G, Mortazavi A, Kwan G, van Baren MJ, Salzberg SL, Wold BJ, Pachter L. 2010. Transcript assembly and quantification by RNA-Seq reveals unannotated transcripts and isoform switching during cell differentiation. Nat. Biotechnol. 28:511–515.

Tsai ZT -Y, Lloyd JP, Shiu S-H. 2017. Defining Functional Genic Regions in the Human Genome through Integration of Biochemical, Evolutionary, and Genetic Evidence. Mol. Biol. Evol.

Tsai ZT -Y, Shiu S -H, Tsai H-K. 2015. Contribution of Sequence Motif, Chromatin State, and DNA Structure Features to Predictive Models of Transcription Factor Binding in Yeast. PLoS Comput. Biol. 11:e1004418.

VanBuren R, Bryant D, Edger PP, Tang H, Burgess D, Challabathula D, Spittle K, Hall R, Gu J, Lyons E, et al. 2015. Single-molecule sequencing of the desiccation-tolerant grass Oropetium thomaeum. Nature 527:508–511.

Wang Z, Zang C, Rosenfeld JA, Schones DE, Barski A, Cuddapah S, Cui K, Roh T -Y, Peng W, Zhang MQ, et al. 2008. Combinatorial patterns of histone acetylations and methylations in the human genome. Nat. Genet. 40:897–903.

Xu S, Grullon S, Ge K, Peng W. 2014. Spatial clustering for identification of ChIP-enriched regions (SICER) to map regions of histone methylation patterns in embryonic stem cells. Methods Mol. Biol. 1150:97–111.

Yamada K, Lim J, Dale JM, Chen H, Shinn P, Palm CJ, Southwick AM, Wu HC, Kim C, Nguyen M, et al. 2003. Empirical analysis of transcriptional activity in the Arabidopsis genome. Science 302:842–846.

Yang L, Takuno S, Waters ER, Gaut BS. 2011. Lowly expressed genes in Arabidopsis thaliana bear the signature of possible pseudogenization by promoter degradation. Mol. Biol. Evol. 28:1193–1203.

Zhang X, Yazaki J, Sundaresan A, Cokus S, Chan SW -L, Chen H, Henderson IR, Shinn P, Pellegrini M, Jacobsen SE, et al. 2006. Genome-wide high-resolution mapping and functional analysis of DNA methylation in arabidopsis. Cell 126:1189–1201.

Zhao Y, Li H, Fang S, Kang Y, Wu W, Hao Y, Li Z, Bu D, Sun N, Zhang MQ, et al. 2016. NONCODE 2016: an informative and valuable data source of long non-coding RNAs. Nucleic Acids Res. 44:D203–-8.

Zou C, Lehti-Shiu MD, Thibaud -Nissen F, Prakash T, Buell CR, Shiu S-H. 2009. Evolutionary and expression signatures of pseudogenes in *Arabidopsis* and rice. Plant Physiol. 151:3–15.

